# *De novo* design of potent CRISPR-Cas13 inhibitors

**DOI:** 10.1101/2024.12.05.626932

**Authors:** Cyntia Taveneau, Her Xiang Chai, Jovita D’Silva, Rebecca S. Bamert, Brooke K. Hayes, Roland W. Calvert, Daniel J. Curwen, Fabian Munder, Lisandra L. Martin, Jeremy J. Barr, Rhys Grinter, Gavin J. Knott

## Abstract

CRISPR-Cas systems are transformative tools for gene editing which can be tuned or controlled by anti-CRISPRs (Acrs) - phage derived inhibitors that regulate CRISPR-Cas activity. However, Acrs that are capable of inhibiting biotechnologically relevant CRISPR systems are relatively rare and challenging to discover. To overcome this limitation, we describe a highly successful, rapid, and generalisable approach that leverages *de novo* protein design to develop new-to-nature proteins for controlling CRISPR-Cas activity. Using CRISPR-Cas13 as a representative example, we demonstrate that AI-designed anti-CRISPRs (AIcrs) are capable of highly potent and specific inhibition of CRISPR-Cas13 proteins. We present a comprehensive workflow for design validation and demonstrate AIcrs functionality in controlling CRISPR-Cas13 activity in bacteria. The ability to design bespoke inhibitors of Cas effectors will contribute to the ongoing development of CRISPR-Cas tools in diverse applications across research, medicine, agriculture, and microbiology.

## MAIN

Clustered regularly interspaced short palindromic repeat (CRISPR) and CRISPR associated (Cas) effectors are microbial adaptive immune systems that carry out interference against mobile genetic elements^1^. CRISPR-Cas systems function through a Cas effector protein (typically nuclease) guided by a CRISPR-RNA (crRNA) to base pair with foreign genetic material. Base pairing between the crRNA and the target nucleic acid triggers a conformational change in the Cas effector that activates the nuclease domain leading to target restriction^2^. CRISPR-Cas systems are categorized by their composition and broader mechanism of interference through a tiered classification of Class (1 or 2), type (I, II, III, IV, V, or VI), and subtype^3^. Of particular interest are the Class 2 CRISPR-Cas effectors (Cas9, Cas12, and Cas13); enzymes that have transformed biotechnology with applications in precision gene editing, transcriptome engineering, and diagnostics^4^. The widespread application of these systems highlights the need for effective tools to selectively control and tune their activity^5,6^.

Anti-CRISPRs (Acrs) are mobile genetic element encoded proteins or RNAs that inhibit the activity of CRISPR-Cas machinery^7,8^. Originally discovered with the observation of resistance to CRISPR-Cas targeting^9^, studies have since leveraged guilt-by-association with known anti-CRISPR associated (*aca*) genes or evidence of self-targeting to uncover potent anti-CRISPRs capable of disarming Cas9^10^, Cas12^11,12^, or Cas13^13,14^. Like the CRISPR-Cas systems they target, Acr nomenclature abides by a tiered classification system of class and subtype (e.g. AcrIIA4 – the fourth inhibitor of Type II-A CRISPR-Cas9). Many Acrs exert their inhibitory effects directly on the Cas effector nuclease by blocking crRNA or target/activator nucleic acid binding sites, or the conserved catalytic centre of the nuclease (**Fig.1**). Once bound, Acrs function through diverse mechanisms including the prevention or disruption of active effector complex formation^15^, crRNA degradation^16^, and competitive or allosteric inhibition^17,18^. This diverse toolbox of inhibitors supports the precision control of gene editing in human cells to limit unintended edits^5^, control gene drives in the environment^19,20^, or as selectable markers in phage engineering^21^. To expand the toolbox of available inhibitors, researchers have turned to engineering, deep-learning, and structure-guided Acr discovery to broaden target range or Acr potency^22–25^. However, the diversity of known CRISPR-Cas effectors far outstrips the number of functionally validated and potent anti-CRISPRs. This is particularly evident for the Type VI RNA-guided, RNA-targeting CRISPR-Cas13 systems for which only three validated inhibitors have been described^13,14,22,26^, leaving many of the biotechnologically relevant Cas13 systems without Acrs.

**Fig. 1.**
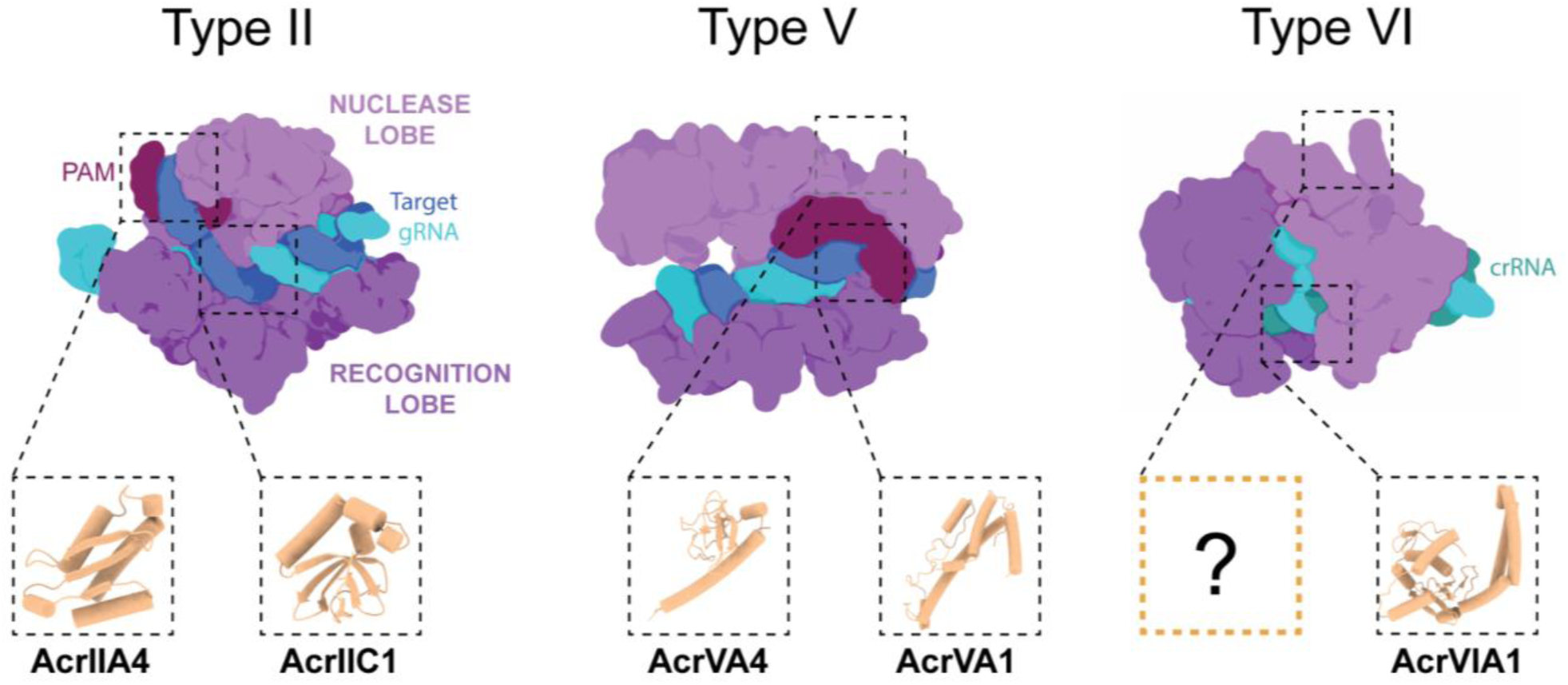
Diverse examples of phage derived anti-CRISPRs (Acrs) targeting Type II CRISPR-Cas9, Type V CRISPR-Cas12a, and Type VI CRISPR-Cas13. Anti-CRISPR (Acr) proteins employ diverse mechanisms to inhibit CRISPR-Cas systems through interactions with the Nuclease or Recognition lobes. Type II (Cas9), AcrIIA4 (PDB: 5XN4^50^) and AcrIIC1 (PDB: 5VGB^33^) block DNA target recognition and the HNH nuclease, respectively. Type V (Cas12a), AcrVA4 (PDB: 6NMA^51^) allosterically inhibits DNA binding and AcrVA1 (PDB: 6NMD^51^) cleaves the crRNA to prevent target DNA recognition. Type VI (Cas13), AcrVIA1 (PDB: 6VRB^13^) inhibits RNA cleavage by blocking activator RNA recognition. No inhibitors have been described that block the highly conserved higher eukaryotic/prokaryotic nucleotide binding (HEPN) domain of Type VI CRISPR-Cas13.

To overcome limitations in the discovery of naturally occurring Acrs, we leveraged *de novo* protein design to create new-to-nature protein inhibitors of CRISPR-Cas which we call AI-designed Acrs (AIcrs). Using unconditional protein generation with RosettaFold-Diffusion (RFdiffusion)^27^ and inverse folding with ProteinMPNN^28^, we describe the design of AIcrs against *Leptotrichia buccalis* (Lbu) CRISPR-Cas13a – a representative CRISPR-Cas13 family member for which no known or validated Acrs exist. We establish a generalisable approach to the design of AIcrs and demonstrate that AIcrs are highly compact, potent, and specific inhibitors of the targeted CRISPR-Cas machinery. Through extensive validation, we show that AIcrs function via a mode of action consistent with their design and demonstrate potent inhibition of CRISPR activity in bacteria. AIcrs have the potential to overcome limitations of traditional discovery-based methods to provide effective and versatile inhibitors that enhance the safety, efficacy, and control of gene editing and beyond.

## RESULTS

### *De novo* design of AIcrs against the conserved Cas13 nuclease domain

Phage derived anti-CRISPRs are highly structurally diverse, and typically exploit core functions in CRISPR-Cas assembly, activation, or catalytic activity. To explore the application of *de novo* protein design in generating potent anti-CRISPRs, we targeted the well-studied and structurally characterised LbuCas13a for which no known natural Acrs exist^29^. To inhibit LbuCas13a activity, we generated designs against the higher eukaryotes and prokaryotes nucleotide-binding (HEPN) endoribonuclease domain, a domain that is strictly conserved across Type VI CRISPR-Cas13 systems. The HEPN nuclease cleaves substrate RNA in response to crRNA hybridization to a complementary activator-RNA (aRNA)^30,31^. Using RFdiffusion, we unconditionally generated protein scaffolds to create AIcrs that would directly compete with substrate RNA recognition via interaction with the conserved HEPN nuclease active site pocket at residue (H473) and proximal surface accessible hydrophobic residues (V411, V421, F995) (**Fig.2a)**. From a set of 10K designs, we selected 96 candidates using a combination of in-house and previously published *in silico* metrics^27^. The final set of 96 AIcr designs were then analysed by multiple structural alignment (MSTA) (**Fig.2b, Supplementary Fig.1**). AIcr designs ranged from 70-127 amino acids in length with highly diverse structures, consistent with unconditional protein fold generation (**Supplementary Fig.2**). A search of the predicted AIcr structures against the Protein Data Bank or AlphaFold Database revealed no significant similarity to known proteins. Examining the MSTA guide tree revealed several mixed αβ designs and numerous related clusters of predominantly α-helical designs (**Fig.2b, Supplementary Fig.2**). Across the structurally distinct clusters, and within structurally related clusters there was no clear sequence conservation (**Supplementary Fig.1**). All AIcrs filtered into the final set exhibited relatively globular shapes (**Supplementary Fig.3)** and were predicted to have negatively charged surface potentials (isoelectric points 3.6 - 4.9) (**Supplementary Fig.4**); consistent with design against a positively charged nucleic acid binding site and akin to many phage-derived Acrs. Examining AlphaFold2 (AF2) predictions from each AIcrs structural class in complex with the target Cas13 suggested diverse mechanisms of target hotspot engagement through charged or hydrophobic interactions, in addition to HEPN active site shape complementarity (**Supplementary Fig.4, 5, 6**).

**Fig. 2.**
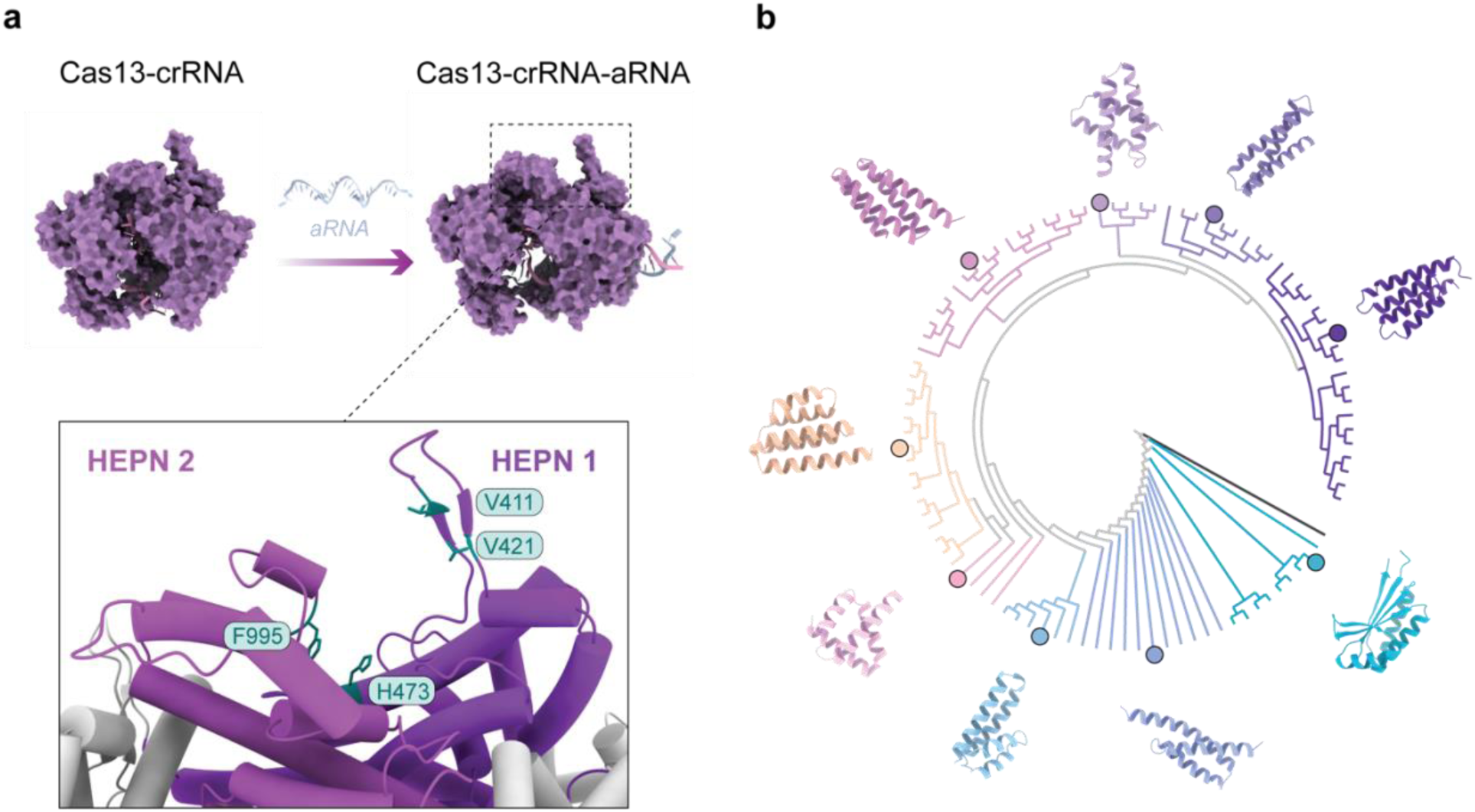
AIcr designs targeted to the LbuCas13a HEPN domain. **a**, Structure of *Leptotrichia buccalis* (Lbu) Cas13a (surface, purple) bound to crRNA (pink) hybridized to an activator-RNA (aRNA, pale blue) drives activation of the HEPN nuclease (boxed region) (PDB codes 5XWY and 5XWP^29^). Design hotspots (stick representation, teal) are shown between HEPN1 (cartoon, dark purple) and HEPN2 domains (cartoon, light purple). **b**, Multiple structure alignment (MSTA) tree for the 96 filtered AIcr designs rooted to GFP (black) and coloured by structural class. Representative structurally diverse AIcrs are shown (cartoons at circled nodes).

### AIcrs are functional inhibitors of CRISPR-Cas13 activity

To rapidly screen each AIcr for the ability to inhibit LbuCas13a HEPN nuclease activity, we developed a cell-free expression and Cas13a activity assay system. Microgram quantities of unpurified cell-free expressed AIcrs were supplemented into an LbuCas13a HEPN nuclease activity assay containing LbuCas13a-crRNA complex, complementary activator RNA (aRNA), and a fluorescence-quenched reporter substrate RNA (**Fig.3a**). In the absence of inhibitors, activated LbuCas13a freely cleaves the reporter RNA resulting in an increase in fluorescence (**Supplementary Fig.7**). In the presence of cell-free expressed AIcrs, we observed a greater than 50% reduction in fluorescence after 60 mins for 10 of the 96 designs, indicative of a reduction in LbuCas13a activity (**Fig.3a, Supplementary Fig.7**). To further investigate AIcr-mediated inhibition, we measured the association of LbuCas13a complexes (crRNA +/- aRNA) to anti-His antibody immobilized AIcrs by biolayer interferometry (**Fig.3b, Supplementary Fig.8-9**). Broadly consistent with the coupled cell-free and Cas13a activity assay, seven AIcrs displayed binding to the activated LbuCas13a-crRNA-aRNA complex and, of these, five displayed binding to the LbuCas13a-crRNA complex (**Supplementary Fig.8-9**).

**Fig. 3.**
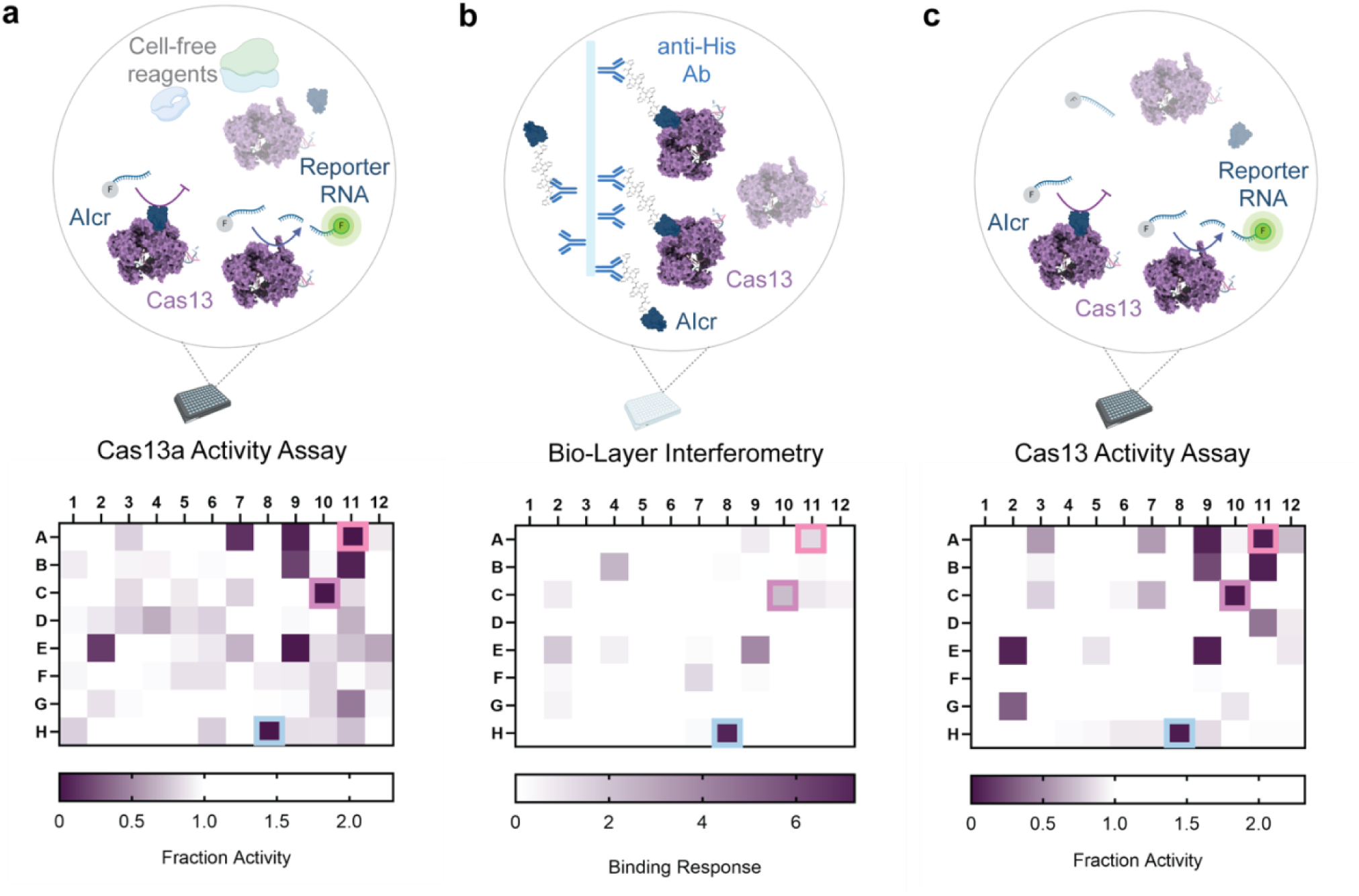
AIcrs are functional inhibitors of LbuCas13a activity. Schematic representations of the experiment (top row) and heat maps corresponding to a 96-well plate layout (bottom row) were each well contains a single designed AIcr. **a,** RNase activity of LbuCas13a-crRNA-aRNA in the presence of cell-free expressed AIcrs. Fluorescence values at 60 mins are shown normalized to LbuCas13a-crRNA-aRNA activity alone. **b,** Biolayer interferometry binding assay measuring the interaction between LbuCas13a complexes and His-AIcrs, immobilized via an anti-His antibody. Maximum binding response height is shown normalized to the AIcr load response. **c**, RNase activity of LbuCas13a-crRNA-aRNA in the presence of purified AIcrs. Normalization and timing as in **a**. Three lead candidates (A11, C10, and H8) were selected for further characterization (boxed).

To validate our initial screen, we partially purified all 96 AIcr designs using batch-based affinity chromatography (**Supplementary Fig.10**) and measured their ability to inhibit LbuCas13a HEPN nuclease activity in a semi-purified system. We observed near perfect agreement between the inhibition activities of AIcrs generated in either the cell-free or semi-purified setting (**Fig.3c, Supplementary Fig.11**). With a success rate of 10% and an 8-week turnaround time from design to validated inhibition, the generation of AIcrs as inhibitors of CRISPR-Cas is highly cost effective, feasible, and presents an efficient alternative to the discovery of naturally encoded Acrs. We selected three AIcrs, A11, C10 H8, that displayed consistent inhibition activity, for further investigation and rigorous validation. In line with convention in the field, and to differentiate them from naturally occurring phage-derived inhibitors, we named the three validated Type VI LbuCas13a inhibitors AIcrVIA1 (A11), AIcrVIA2 (C10), and AIcrVIA3 (H8).

### Cas13a AIcrVIAs are potent, highly stable, and true to design

To investigate the potency of AIcrVIA1 – VIA3, we titrated each purified AIcrVIA against a fixed concentration of the LbuCas13a-crRNA-aRNA complex and measured the velocity of the activated HEPN nuclease (**Fig.4a**). Calculating the concentration of AIcr required to achieve 50% inhibition (IC_50_) of LbuCas13a revealed values in the low nanomolar range for all three AIcrVIAs, consistent with high affinity binding. To validate that the AIcrs reflected the overall structure of the designs, we measured the secondary structure composition of each AIcrVIA using circular dichroism (CD) (**Fig.4b**). CD spectra for the purified AIcrVIAs revealed secondary structural fingerprints that reflect the designs with evidence for the mixed αβ and loop composition of AIcrVIA1, the exclusively α-helical composition of AIcrVIA2, and the predominantly β-stranded composition of AIcrVIA3. Moreover, consistent with other AI-designed proteins and the naturally occurring Acrs, AIcrVIAs were highly robust and thermostable (**Fig.4b**). Having purified all AIcrVIAs to homogeneity using size exclusion chromatography, we observed that each AIcrVIA eluted at a volume broadly with the designed monomeric form (**Supplementary Fig.12**).

**Fig. 4.**
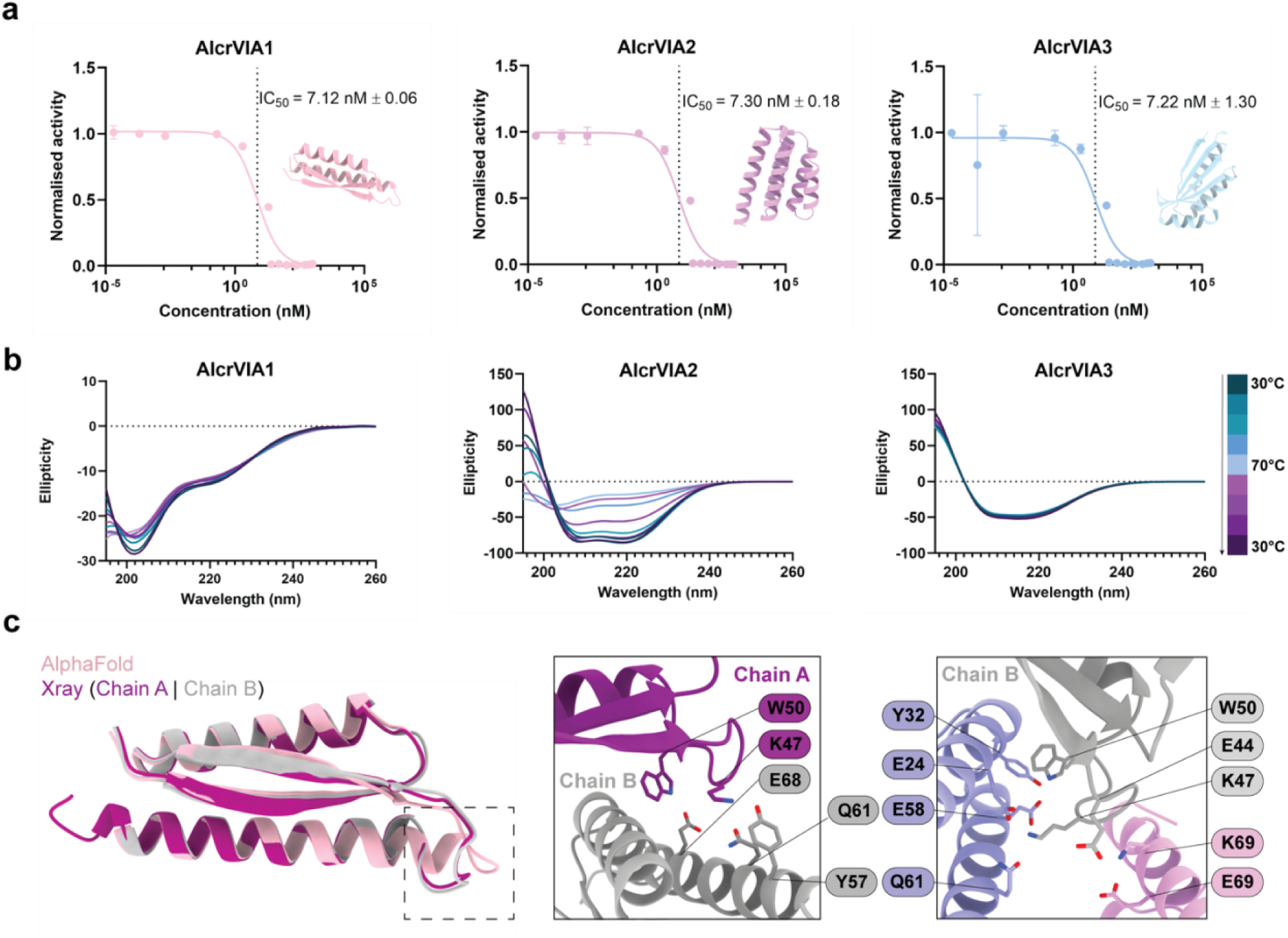
AIcrVIAs are potent inhibitors and true to design. **a,** Inhibitor dose dependence of LbuCas13a HEPN nuclease activity for AIcrVIA1 (left, pink), AIcrVIA2 (middle, dark pink), and AIcrVIA3 (right, blue). IC₅₀ values (∼7 nM) indicate high inhibitory potency. Data are shown as mean ± s.d. (*n* = 3). Cartoon representations for each AIcr design are shown. **b,** Circular dichroism (CD) analysis of AIcrs fold and stability across a temperature range. CD spectra of AIcrVIA1 (left), AIcrVIA2 (middle), and AIcrVIA3 (right) were recorded from 30°C to 70°C (blue gradient, heating) and back to 30°C (purple gradient, cooling) in 10°C increments. The ellipticity curves reflect thermal stability and refolding behaviour**. c,** X-ray crystal structure of AIcrVIA1 (cartoon, magenta and grey) superposed to the AF2 prediction (cartoon, pink) with the variable loop highlighted (dashed box). The interaction interface for the variable loop is shown for each copy of within the asymmetric unit (right panels).

To further validate congruence between design and reality, we determined the X-ray crystal structure of AIcrVIA1 to 1.9 Å resolution (**Fig.4c, Supplementary Table 1**). AIcrVIA1 crystallised with two copies in the asymmetric unit, each sharing a βαβα topology and an overall root mean square deviation (rmsd) of 0.64 Å (chain A to B). Consistent with the AF2 prediction, both copies of AIcrVIA1 resembled the designed structure (rmsd of 1.7 and 1.8 Å, His-tag excluded for chain A and B, respectively). The only deviations between the AF2 model and the experimental structure resulted from extensive crystal contacts in residues 47–52 (**Fig.4c**). Taken together, these biophysical and structural data provide clear evidence that AIcrVIAs are highly potent, stable, and their structures reflect their intended design.

### AIcrVIAs competitively inhibit the HEPN nuclease

AIcrVIA1 – AIcrVIA3 were designed to competitively inhibit the LbuCas13a nuclease via direct association with the conserved HEPN. To demonstrate this mode of inhibition, we first assayed aRNA binding to the LbuCas13a-crRNA complex using fluorescence anisotropy in the presence or absence of an excess of each AIcr (**Fig.5a**). With an excess of AIcrVIA in competition with the crRNA and aRNA, binding to a complementary aRNA was unperturbed suggesting the absence of any non-specific associations that would compete with crRNA or aRNA association. We next validated that each AIcrVIA was capable of a direct interaction with the activated LbuCas13a-crRNA-aRNA complex by size exclusion chromatography. Consistent with our biolayer interferometry binding data, each AIcrVIA stably eluted with the activated LbuCas13a complex (**Supplementary Fig.12**) and the presence of any AIcrVIA stabilized both the LbuCas13a-crRNA or LbuCas13a-crRNA-aRNA complex (**Fig.5b**).

**Fig. 5.**
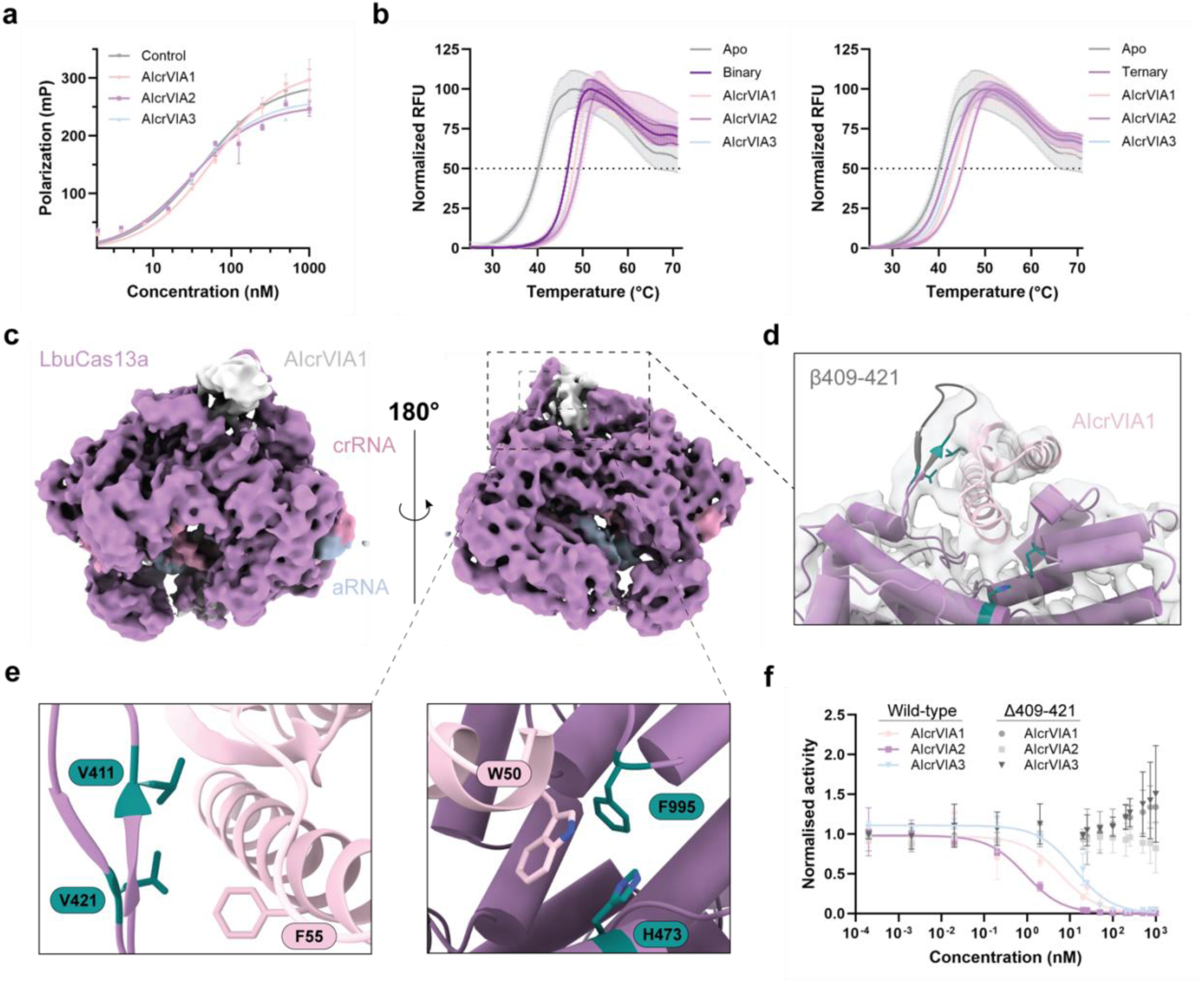
AIcrVIAs are competitive inhibitors of the LbuCas13a HEPN nuclease. **a,** Fluorescence anisotropy (polarization) assay assessing aRNA binding to LbuCas13a-crRNA in the presence or absence (control) of an excess of AIcrVIAs (*n* = 3 with mean ± s.d.). **b,** Melt curve analysis of LbuCas13a-crRNA (left) and LbuCas13a-crRNA-aRNA with and without AIcrVIs. RFU, relative fluorescence units; Apo, LbuCas13a; Binary, LbuCas13a-crRNA; Ternary, LbuCas13a-crRNA-aRNA(*n* = 33 with mean ± s.d.). **c,** Cryo-EM map of the LbuCas13a ternary complex with AIcrVIA1, coloured by nucleic acid (crRNA in pink, aRNA in blue) and protein (LbuCas13a in purple). Unassigned density is attributed to AIcrVIA1 (grey). **d,** Cryo-EM structure highlighting the interaction of AIcrVIA1 (pink) with the domains (purple) and the ß-turn loop (residues 409–421, black). Hotspots residues designated at the design stage are shown as stick (teal). **e,** LbuCas13a hotspots (sticks, teal) interacting with AIcrVIA1 aromatic residues (F55 and W50). **f,** LbuCas13a HEPN nuclease activity normalised to no-inhibitor comparing wild-type or LbuΔ409–421 in the presence of increasing concentrations of AIcrVIAs. Data are shown as mean ± s.d. (*n* = 3).

To visualize an AIcrVIA in the act of inhibiting LbuCas13a, we determined a 3.55 Å cryo-EM structure of the activated wild-type LbuCas13a complex in the presence of AIcrVIA1 (**Fig.5c, Supplementary Fig.13-14**). The overall structure of the activated LbuCas13a within the complex resembled that of the previously published X-ray crystal structure with an overall rmsd of 1.4 Å (PDB 5XWP, **Supplementary Fig.15a**). We observed clearly resolved secondary structural features for AIcrVIA1 within the HEPN nuclease, albeit with limited high-resolution information due to local conformational heterogeneity driven by continuous shifts in AIcrVIA1 around the HEPN nuclease (**Supplementary Video 1**). Despite the apparent dynamics, the resolved cryo-EM structure closely matched the final prediction for the complex LbuCas13a-AIcrVIA1 obtained from the AI design workflow (AF2 initial guess implemented in MPNN) – with an overall rmsd of 1.1 Å (**Supplementary Fig.15b-c**). AIcrVIA1 is positioned within the HEPN nuclease making numerous contacts with structural elements proximal to the conserved HEPN nuclease, including a highly positively charged β-stranded element (residues 409-421) (**Fig.5d, Supplementary Fig.16a-b**). Closer examination of the interaction interface revealed regions of AIcrVIA1 near the designated HEPN nuclease hotspots, illustrative of a congruence between design and experimental reality (**Fig.5e, Supplementary Fig.16c**).

Given the observed extensive contacts between AIcrVIA1 and the LbuCas13a β-stranded element, we generated a deletion construct lacking β-stranded element (LbuΔ409-421) to validate the interactions. Assaying LbuΔ409-421 HEPN nuclease activity in the presence of AIcrVIA1 revealed a completely abolished inhibition efficacy, supporting the observed interactions (**Fig.5f**). Furthermore, the inhibition activity of both AIcrVIA2 and AIcrVIA3 were completely abolished against LbuΔ409-421, suggesting that these AIcrs likely occupy the same HEPN nuclease pocket, consistent with their design against the same features and hotspots (**Fig.5f**). Taken together, these findings point to a highly specific mechanism of HEPN nuclease competitive inhibition for all three AIcrVIAs as designed.

### AIcrs are specific and control Cas13 RNA editing in bacterial cells

Phage-derived Acrs are highly successful tools to control the activity of CRISPR-Cas systems in bacterial and human cells^5,6^. Critical to Acr applications is specificity where many phage-derived Acrs are highly specific to their target Cas effector with only a few examples of broad spectrum Acrs^16,32,33^. Given the validated interaction of AIcrs with the LbuCas13a HEPN nuclease, we hypothesised that the AIcrsVIAs would show limited inhibition of related, non-Lbu CRISPR-Cas13 effectors. To investigate their specificity, we assayed the three AIcrVIAs against the functionally and evolutionarily related *Lachnospiraceae bacterium* (Lba) Cas13a, *Thermoclostridia caenicola* (TccCas13a) Cas13a, and *Ruminococcus flavefaciens* (Rfx) Cas13d. Concentration titrations of AIcrVIAs in far excess of active Cas13 enzyme revealed no apparent inhibition of any Cas13 tested, validating the high specificity of the designs (**Fig.6a, Supplementary Fig.17**). Combined with their validated potency, the observed specificity underscores the precision of AIcrVIAs and supports their potential for targeted inhibition in biotechnological applications.

**Fig. 6:**
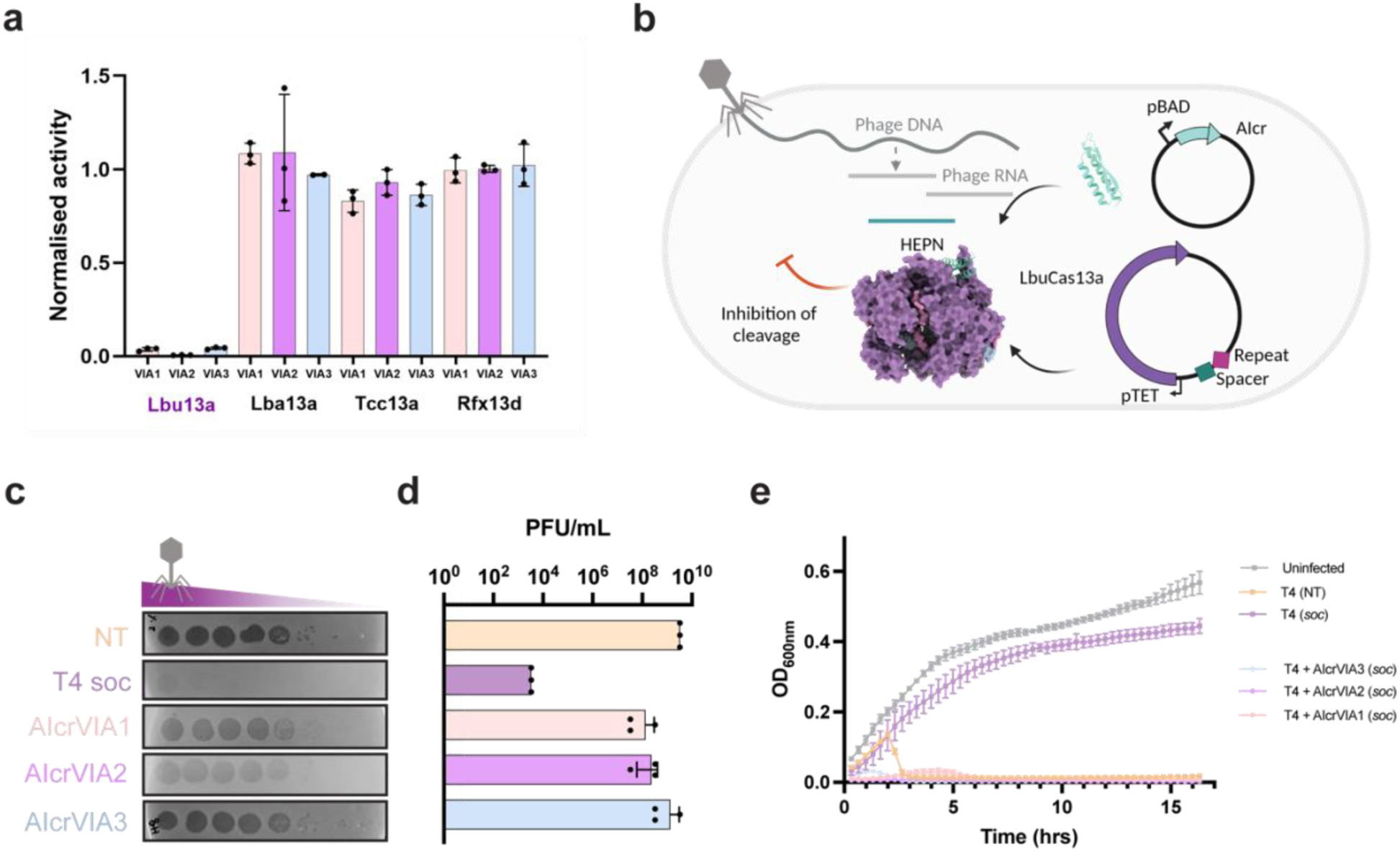
AIcrs are specific and inhibit Cas13 in bacterial cells. **a,** Activity of various Cas13 effectors in presence of a 10-fold excess of AIcrVIA (*n* = 3). Lbu13a = LbuCas13a; Lba13a = LbaCas13a; Tcc13a = TccCas13a; Rfx13d = RfxCas13d; VIA1 = AcrVIA1; VIA2 = AIcrVIA2; VIA3 = AIcrVIA3. **b,** Schematic of the phage assay to assess AIcrVIA efficacy in *Escherichia coli* co-expressing LbuCas13a and crRNA. Induction of LbuCas13a enables defence against T4 phages through non-specific RNA degradation upon phage genome recognition. Induction of AIcr inhibits LbuCas13a activity to allow T4 replication. **c,** Phage plaque assay performed on *E. coli* co-expressing LbuCas13a and crRNA targeting T4-*soc*, incubated with serial dilutions of T4 phage (decreasing concentration from left to right). Results were quantified in **d**, with efficiency of plaquing (EOP) represented as PFU/mL for each AIcrVIA. Inducible expression of AIcrVIAs restores T4 phage titres across induction levels, indicating potent inhibition of LbuCas13a activity in *E. coli*. **e,** Phage liquid culture assay demonstrating reduction in *E. coli* growth in the presence of AIcrVIAs, confirming inhibition of LbuCas13a in vivo. Data from **d** and **e** are shown as mean ± s.d. (*n* = 3 independent experiments). NT = non-targeting crRNA; T4 soc = T4 soc-targeting crRNA; PFU = plaque-forming units.

To explore the applicability of AIcrVIAs in bacterial cells, we first constructed an anhydrous tetracycline (aTc) inducible LbuCas13a vector with a constitutively expressed non-targeting (NT) or T4 *soc-*targeting (T) crRNA (**Fig.6b**). *E. coli* expressing LbuCas13a and T4 *soc-*targeting crRNA effectively restricted T4 phage replication, whereas non-targeting crRNA had no effect (**Fig.6c-d**) – consistent with the RNA-guided HEPN nuclease activity of CRISPR-Cas13. Next, we constructed an arabinose inducible AIcrVIA expression vector which was co-transformed with the LbuCas13a-crRNA T or NT vector (**Fig.6b**). In contrast to the conditions lacking AIcr, the inducible expression of either AIcrVIA1, VIA2, or VIA3 recovered T4 phage titres across a range of induction strengths, demonstrating potent inhibition of LbuCas13a activity in *E. coli* (**Fig.6c-d, Supplementary Fig.18-19**). Examining the effect of AIcrVIAs in phage liquid culture assays using the same inducible system revealed a drastic reduction of *E. coli* growth in the presence of T4 phage and AIcrVIAs (**Fig.6e**). These data demonstrate that the AIcrVIAs fulfil the same biological function as natural phage derived Acrs and provide a strong indication for their applied potential in phage engineering.

## DISCUSSION

Anti-CRISPR (Acr) proteins remain crucial for the post-translational regulation and control of CRISPR-Cas systems in gene editing applications. To date, and after over ten years of research^34^, there are 118 experimentally validated Acrs targeting a diverse array of CRISPR-Cas systems, many of which are not often utilised in biotechnology^35^. The hunt for naturally occurring inhibitors continues to be challenging, even with cutting-edge developments in deep-learning and structure-guided discovery^22–25^. Here we implemented a rapid approach to Acr design that leverages AI-driven unconditional protein fold generation to *de novo* create compact, potent, and highly specific AI-designed anti-CRISPRs (AIcrs). This approach is incredibly rapid, taking eight weeks from target selection to verified nanomolar inhibition of CRISPR-Cas activity – significantly faster than traditional protein discovery workflows. Using this approach, we designed first in class AIcrVIAs targeted to the HEPN nuclease of LbuCas13a, a CRISPR-Cas system utilized for phage engineering and molecular diagnostics with no known Acrs^36–38^. The development of potent AIcrVIAs provides selectable markers for phage engineering applications, akin to the application of phage derived AcrVIA1 for phage engineering with *L. seeligeri* Cas13^21^. By leveraging targeted inhibitors, it may also be possible to fine-tune the interactions between phages and bacterial hosts, potentially leading to more effective treatments for bacterial infections. This approach may not only enhance the therapeutic potential of phages but also opens new avenues for research and development in the field of synthetic biology. Interestingly, like many naturally occurring Acrs^6^, our AIcrs exhibited no significant structural similarity to known proteins, a fact that stifled attempts to structurally bootstrap towards naturally encoded Acrs. While the designs were not predictive of natural occurring Type VI Acrs, their ability to inhibit LbuCas13a *in vivo* suggests the possibility that phage may have evolved HEPN inhibitors. With a larger sample size of AI-designed proteins to extract key interaction interfaces, it may be possible to discover Acrs similar to recent work identifying plant and fungal pathogen interactions^39^. Overall, our application of *de novo* protein design to generate AI-designed anti-CRISPRs (AIcrs) represents a powerful approach to overcoming the limitations of traditional discovery-based approaches.

While the discovery of naturally encoded Acrs yields mechanistically diverse inhibitors, approaches leveraging AI-protein design afford user-defined control over the mode of inhibition (crRNA/DNA/RNA competition, active site inhibition, allosteric inhibition). With an understanding of functionally relevant target sites, AI-driven protein design approaches could be applied to design bespoke inhibitors of any CRISPR-Cas Class, Type, or subtype, including those for which no Acr is known to exist. For example, AIcrs could be designed to inhibit CRISPR Cas1-Cas2 acquisition machinery to develop switchable molecular recorders^40^ or tailored to regulate emerging classes of editors including TnpB, IsrB, IscB, Fanzor, as well as the numerous CRISPR-Cas guided DNA integrases^41–43^. A very recent study described the development of BindCraft, a novel AI-driven approach to protein design with a demonstrated capability to design inhibitors of *S. pyogenes* Cas9 with applications in controlling its gene editing activity^44^.

Beyond CRISPR-Cas, the advent of AI-driven protein design (e.g. RF-diffusion) has significantly transformed *in silico* protein design, dramatically improving the efficiency of the design process and the success rate^27^. While revolutionary, the field is in its infancy and rapidly developing. At present AI-protein design outcomes remain difficult to predict. From our set of 96 AIcrs we recovered 10 efficient inhibitors for which no single *in silico* parameter was predictive of success *in vitro* (**Supplementary Fig.20**). Retraining on the increasing volumes of wet-lab data will pave the way for a higher success rate in future, especially with the use of high-throughput quantitative cell-based screens. The development of one-shot protein design methods offers a tantalising potential to overcome the need for screening a subset of designs^44^. However, target protein flexibility and conformational dynamics remain a significant challenge, especially in cases where structural insights may be limited. Furthermore, protein targets rarely exist in isolation, but significant advances in nucleic acid and co-factor interaction prediction using AlphFold3^45^, Boltzmann generative models like Boltz-1^46^, or Chai-1^47^ may overcome this challenge. Additionally, tools such as RoseTTAFold All-Atom^48^ show promise for designing or targeting these multi-component, heterogeneous complexes. Likely, networks will need retraining on highly curated datasets specific to preserve unique structural and functional properties of protein families. This approach contrasts with models trained on the PDB, which excel at generating robust proteins but may miss the nuanced features essential for the specificity and function of protein families. Tailored protein design will likely require a balance between the robustness provided by access to large training datasets, and the precision afforded by training datasets tailored to specific protein families, functions or properties^49^. As the field rapidly accelerates it seems inevitable that AI-driven protein design will have far reaching impact across biological research.

## Methods

### Computational design and analysis of AIcrs

The design of specific proteins targeting the HEPN nuclease of *Leptotrichia buccalis* (Lbu) Cas13a was carried out using RFdiffusion^27^. A total of 10K 70-130 amino acid designs of α and mixed αꞵ topology were generated unconditionally against hotspots V411, V421, H473, and F995. Inverse folding was conducted using ProteinMPNN^28^, followed by energy minimization using FastRelax. Prediction quality was assessed by AlphaFold2 initial guess implemented in ProteinMPNN. Designs were filtered by predicted aligned error (PAE) interaction scores <10 ^52^. This subset of 80 designs underwent further rounds of ProteinMPNN (iterations or serial) to minimize PAE interact scores. The final set of 96 candidates were selected based on examining AlphaFold2 initial guess output models for favourable design to target interactions, overall pLDDT, and target root mean square deviation (rmsd). AI design diversity was assessed by multiple structure alignment (MSTA) using FoldMason^53^ with GFP root (default parameters - gap open penalty 10, gap extension penalty 1). The MSTA was visualised and annotated using iTol^54,55^. Multiple sequence alignments were visualised using JalView^56^. Parameters describing design and target interactions (interface area (Å^2^), ΔG, Dmax, asphericity, radius of hydration) were calculated using the Python package MDAnalysis^57^ and PISA^58^. Each AIcr was *E. coli* codon-optimized and cloned into pET-29(+) with an in-frame C-terminal hexahistidine tag (Twist Bioscience).

### Expression and purification of Cas13 proteins

Cas13 homologs and their sequences are listed in **Supplementary Table 3**. Each recombinant Cas13 was expressed as previously described^59^. Briefly, each Cas13 was expressed in Terrific broth (TB) media (with appropriate antibiotic) seeded with 0.5% (v/v) of overnight culture, incubated at 37℃ at 180 RPM until of 0.5-0.6, cold shocked for 20 mins and induced with 0.5 mM IPTG followed by incubation at 16℃ 180 rpm for 16 hrs. Cells were harvested by centrifugation (4,000xg for 20 mins) and flash frozen in liquid nitrogen prior to storage at −80℃. All buffers used to purify Cas13 homologs were the same except for the pH for TccCas13a (pH7.5 throughout). Cells were resuspended in lysis buffer (50 mM Tris pH 7.0, 500 mM NaCl, 10 mM Imidazole, 5% (v/v) glycerol, 1 mM TCEP, 0.5 mM PMSF, EDTA-free protease inhibitor cocktail (Roche)), and lysed by sonication. The lysate was clarified by centrifugation (35,000xg, 45 mins at 4℃) and the supernatant incubated with Ni-NTA resin (Qiagen) for 1 hour at 4℃. The resin was washed with 10 CV (column volume) of buffer 1 (20 mM Tris pH 7.0, 500 mM KCl, 10 mM Imidazole, 5% (v/v) glycerol, 1mM TCEP), followed by 10 CV of buffer 1 supplemented with 20 mM Imidazole, followed by elution with 5 CV of buffer 1 supplemented with 300 mM Imidazole. Cas13 fractions were pooled and dialysed overnight at 4℃ (14K MWCO) against 50 mM HEPES, 150 mM NaCl, 5% (v/v) glycerol, 1 mM TCEP in the presence of 1:50 mg weight ratio of TEV protease if MBP tagged. The filtered dialysate was loaded onto a 5mL HiTrap SP (Cytiva) and eluted over a 10 CV KCl gradient (0.15-1M). Peak fractions were pooled and concentrated (30K MWCO, Amicon) prior to size exclusion chromatography (S200 16/600, Cytiva) developed in 20 mM HEPES pH 7.0, 200 mM KCl, 10% (v/v) glycerol, 1 mM TCEP. Protein purity was assessed by SDS-PAGE and peak fractions concentrated to 60 µM (30K MWCO, Amicon). Purified protein was snap frozen with liquid nitrogen in aliquots and stored at −80℃.

### Cell-free AIcr production, screening, and data analysis

Cas13a Alcr open reading frames were amplified with T7 forward and reverse primers (**Supplementary Table 3**) using 2X Q5 polymerase Master Mix (NEB) and purified with QiaQuick PCR clean up (Qiagen). Each AIcr was expressed from 2 nM of purified PCR product in PUREFrex 2.0 (GeneFrontier) cell-free expression system prepared in accordance with the manufacturer’s protocol. The reaction mixture was incubated for 4 hours at 37°C and held at 4°C until assaying. LbuCas13a-crRNA ribonucleoprotein (RNP) complex (400 nM LbuCas13a and 400 nM crRNA (Integrated DNA Technologies, IDT)) was prepared in 1X Cas13 cleavage buffer (20 mM HEPES-K pH 6.8, 50 mM KC, 5 mM MgCl_2_ and 5% (v/v) glycerol) at room temperature for at least 15 mins and no longer than one hour. AIcr and substrate reporter mix was prepared as 10 µL containing 0.67 µM fluorescence reporter RNA, 13.33 pM aRNA in 3X Cas13 cleavage buffer prior to the addition of 20 µl in vitro translated AIcr solution. Cas13a HEPN nuclease activity was assayed in black 384-well plates (Corning Cat#3820) by mixing 5 µL of RNP solution with 15 µL of AIcr-aRNA reporter solution. Fluorescence was immediately monitored every 1 min for 2 hrs at 37°C (ClarioSTAR plus, BMG). The fluorescence values at 60 mins were extracted and normalized against no AIcrVIAs controls to express as a fraction of activated LbuCas13a activity (heat map).

### Cas13a-AIcr binding assays

Initial binding assessments were made via bilayer interferometry on a Gator Prime BLI instrument (Solve Scientific), using immobilised AIcrs to assess binding to recombinant LbuCas13 at 30 °C and 1000 rpm shaking throughout assay duration. LbuCas13a complexes were prepared at 5 µM with crRNA alone (1:1.2 molar ratio) or crRNA and aRNA (1:1.2:1.4 molar ration) in 20 mM Tris-HCl (pH 7.5), 150 mM KCl, 5% (v/v) glycerol, 1 mM TCEP, and incubated for 10 mins at room temperature, prior to dilution to 200 nM in Gator buffer Q (PBS pH 7.4, 0.2%(w/v) BSA, 0.02% (v/v) Tween 20). AIcrs from small scale batch purification were diluted 1/20 in buffer Q. AIcrs were immobilised on Anti-His (HIS) Probes for 3 mins, washed with buffer Q for 2 mins, associated with prepared 200 nM Cas13 RNP complex for 3 mins, and dissociated in buffer Q for a further 3 mins. All binding curves were baselined against ‘no AIcr load’ runs. Max height of binding response normalised to AIcr load height response (all in nm) was used to generate the heat map.

### AIcr small scale protein expression and purification

AIcr plasmids were transformed into BL21 (DE3) *E. coli* and single colonies used to inoculate 2 mL cultures in Overnight Express Instant TB media (Novagen) containing 50 µg/mL Kanamycin (Merck). Cultures were grown at 37°C for 8 hrs, followed by 16°C for 16 hrs, centrifuged and cell pellets frozen at −80°C. Thawed cell pellets were chemically lysed (BPer, ThermoFisher Scientific) for 20 mins at room temperature, and clarified by centrifugation at 17,000 xg for 10 mins at 4°C. Supernatant containing soluble AIcrs was incubated with Ni-NTA superflow resin (QIAGEN) for one hour at 4°C, washed twice with buffer A (50 mM Tris pH 7.0, 500 mM NaCl, 5% (v/v) Glycerol, 1 mM TCEP) containing 10 mM and 20 mM imidazole respectively and proteins eluted in buffer A containing 300 mM Imidazole. Following batch purification, concentration and purity of eluted AIcrs was confirmed using Coomassie blue stained SDS-PAGE analysis and Quick Start Bradford (BioRad) assays.

### Cas13a-AIcrVIA competition assays and data analysis

In a bulk reaction, LbuCas13-crRNA RNP complex was prepared by complexing 133 nM LbuCas13a and crRNA (1:1) in 1X Cas13a cleavage buffer at room temperature for 15 mins. The reaction was commenced by addition of 15 μL RNP complex into black 384-well plate with 5 uL of aRNA reporter binder mix (prepared as above, but with purified AIcrs) to give a final concentration of 1 μM AIcr Cas13a candidates (small scale Ni-NTA purified), 100 nM RNP complex, 0.2 μM fluorescence reporter, 10 pM aRNA, 1X Cas13a cleavage buffer. The fluorescence output was measured on a plate reader (ClarioSTAR plus, BMG) for 2 hours every 1 min at 37°C.

For Cas13 AIcrVIA competition assays, 66.66 nM Cas13a RNP complex was assembled by mixing a 2:1 ratio of Cas13:crRNA at room temperature for 15 mins in 1X Cas13a cleavage buffer. In a black 384-well plate, 15 μL of RNP complex was added to 5 uL aRNA reporter binder mix to give a final concentration of 50 nM RNP complex, 1 μM - 0.2 pM AIcr Cas13a candidates, 500 nM fluorescence reporter, and various aRNA concentration for different Cas13 homologs (LbuCas13a - 20 pM, LbaCas13a - 0.5 nM, TccCas13a 1 nM, RfxCas13d - 50 nM, LbuCas13aΔ409-421 - 50 pM) in 1X Cas13a cleavage buffer. The fluorescence output was measured on a plate reader every 30 sec or 1 min for 30 mins at 37°C. Using PRISM v10.1, the IC_50_ was calculated by measuring the linear slope of each reaction from 0-30 mins for LbuCas13a, TccCas13a, RfxCas13d and 0-15 minutes for LbuCas13aΔ409-421 and LbaCas13a. The slopes were analysed with “inhibitor vs response [three parameters]” with constraints for *bottom* set to constant equal to 0, to generate an IC_50_ value.

### Large scale expression and purification of AIcrVIAs

AIcrVIA1-VIA3 sequences are listed in **Supplementary Table 2**. *E. coli* BL21 (DE3) cells were transformed and cultured in TB medium supplemented with kanamycin (50 µg/mL) at 37 °C, 200 rpm. At OD₆₀₀ ∼0.6, cultures underwent a cold shock, and protein expression induced with IPTG to a final concentration of 0.5 mM and subsequently incubated overnight at 18 °C. Cells were harvested by centrifugation at 4,000 xg for 25 mins at 4 °C, washed with lysis buffer (20 mM Tris, pH 7.5; 500 mM NaCl; 10% (v/v) glycerol; 10 mM imidazole), snap-frozen, and stored at −80 °C. Cell pellets were resuspended in lysis buffer with 1 mM TCEP and EDTA-free protease inhibitor, at a ratio of 10 mL per 1 g of wet cell paste. Cells were lysed by sonication and lysates clarified by centrifugation at 40,000 xg for 30 mins at 4 °C. The clarified lysate was applied to Ni-NTA resin and incubated for one hour at 4 °C with gentle agitation. The resin was subsequently washed with 10 CV of lysis buffer with 20 mM imidazole,1 mM TCEP before elution with 5 CV of lysis buffer containing 200 mM imidazole, 1 mM TCEP. Fractions containing the protein of interest were pooled and concentrated using a 3 kDa cutoff concentrator (Amicon) prior to loading on an S75 10/300 column (Cytiva) running in 20 mM Tris (pH 7.5); 150 mM KCl; 5% (v/v) glycerol; 1 mM TCEP. Eluted protein fractions were either snap-frozen in liquid nitrogen or further concentrated using and stored at −80 °C until further purification.

### Fluorescence polarization

LbuCas13a:crRNA RNP complexes were assembled in 1X Cas13a Cleavage buffer for 15 mins at room temperature. RNP complexes were serially diluted 2-fold and incubated with 40 nM 5’6-FAM labelled aRNA and 2 μM AIcrVIAs (final concentration) for two hours at 37°C. The fluorescence polarization and anisotropy values were measured every 5 mins to determine if the system was at equilibrium (ClarioStar plus, BMG LabTech). Fluorescence anisotropy and polarization values were calculated using MARS Data analysis (BMG LabTech) and the curve was fitted using One Site - Specific Binding (PRISM v10.4).

### Circular dichroism

CD spectra were recorded using a J-810 Circular Dichroism Spectrometer (JASCO). Protein samples at a concentration of 0.3 mg/mL were prepared by dilution in water, resulting in a final buffer composition of 0.3 mM Tris pH 7.5, 2.5 mM KCl, and 0.08% (v/v) glycerol. Triplicate wavelength scans were recorded from 260 to 190 nm at 30°C. For thermal denaturation experiments, protein samples (0.3 mg/mL) were heated from 30°C to 70°C to 30°C. Wavelength scans from 260 to 190 nm were recorded at 10°C intervals, following a 5 min incubation period at each target temperature. The recorded data were averaged, baseline corrected and smoothed using JASCO spectra manager (v2). Data were further analysed on GraphPad Prism (v10.4) using the data recorded between 195 and 260 nm.

### Crystallization and structure determination of AIcrVIA1

AIcrVIA1 was crystallized using the hanging-drop vapor diffusion method at 10mg/ml in 0.1 M Bis-Tris propane (pH 7.2) and 2.6 M ammonium sulfate [(NH₄)₂SO₄] at room temperature. A mixture of crystal morphologies (rectangular or stacked plates) were harvested 10 days after crystallisation setup. Crystals were flash frozen directly into liquid nitrogen without cryoprotectant. X-ray diffraction data were recorded on a Dectris Eiger 16M detector at the MX2 beamline of the Australian Synchrotron. Data reduction was performed using XDS^60^. The initial atomic model was obtained by molecular replacement using the AlphaFold2^61^ predicted structure of the AIcrVIA1 polypeptide in Phenix 1.20.1^62^. The model was manually adjusted in WinCoot 0.9.8.93^63^ to fit the electron density map and iteratively refined using phenix.refine.

### Cryo-EM sample preparation and data collection

The LbuCas13-crRNA-aRNA-AIcrVIA1 complex was assembled in cryo-EM buffer (20 mM Tris (pH 7.0), 30 mM KCl, 5 mM MgCl₂, and 2% (v/v) glycerol). LbuCas13a was combined with crRNA, activator RNA, and AIcrVIA in molar ratios of 1:1.2:1.4:5, respectively. Assembly was carried out sequentially with 5 mins incubation at room temperature after each addition to facilitate complex formation. Subsequently, 100 µL of the assembled complex solution was purified and fractionated over a S200 Increase 3.2/300 SEC column (in cryo-EM buffer) attached to an AKTAMicro (Cytiva). The purity and composition of fractions were verified using SDS-PAGE and 12% (w/v) urea polyacrylamide gel electrophoresis. The resulting complex was immediately used to prepare grids for cryo-EM analysis by applying 3.5 µL onto amylamine glow discharged UltrAufoil R1.2/1.3, 300-mesh gold grids (Quantifoil) before blotting with a force of −3 for 3 sec under 100% humidity at 4C (Vitrobot Mark IV, Thermo Fisher Scientific). Grids were immediately plunge-frozen in liquid ethane and stored in liquid nitrogen until imaging. Cryo-EM data were collected using a Talos transmission electron microscope (Thermo Fisher Scientific) operating at 200 kV. A total of 2,544 movies were recorded at a calibrated pixel size of 0.93 Å²/pixel over 50 frames with an exposure of 1 e⁻/Å² per frame, resulting in a total electron dose of 50 e⁻/Å². The defocus range was set between −0.8 and −1.4 µm to optimize image contrast and resolution.

### Cryo-EM data processing and model building

Dose-fractionated frames were corrected for beam-induced motion and radiation damage using MotionCor2 1.4.5^64^. The aligned, dose-weighted averages were then imported into cryoSPARC (v4.5.2) for further processing. Contrast transfer function (CTF) parameters were estimated with CTFFIND 4.1.14^65^. Micrographs were curated at a resolution of 5 Å or higher. Template picking was prepared by performing blob picking in cryoSPARC 4.5.2^66^, followed by five rounds of 2D classification and heterogeneous refinement. Particles selected from the highest-quality heterogeneous refinement density map were used to train a Topaz 0.2.5 model. All particles picked using Topaz 0.2.5^67,68^ were extracted in a 256×256-pixel box before heterogeneous refinement and *ab initio* reconstruction. The resulting particle set was submitted to three rounds of iterative 3D classification and refined using cryoSPARC non-uniform refinement^69^. Conformational flexibility and heterogeneity in the final particle stack (20,357 particles) were investigated using cryoSPARC 3D variability analysis^70^ (resolution filter of 8 Å, three components). The series with the most extensive and continuous movement of AIcrVIA1 was visualised using ChimeraX over 20 frames.

### Model Building

The structure of the LbuCas13a-crRNA-aRNA-AIcrVIA1 complex was built from the previously published LbuCas13a-crRNA-aRNA structure (PDB 5XWP) and the X-ray crystal structure of AIcrVIA1 described in this study. The initial model was fit to the density map in ChimeraX before interative phenix.refine^62^ (v1.18.2, rigid body, minimization global, and adp refinement restrained with secondary structure and AIcrVIA1 reference model restraints) and real space refinement in Coot (v0.9.4^63^). All figures and visualizations were generated using ChimeraX.

### Phage propagation and phage restriction assays

T4 phage were propagated in *Escherichia coli* B strains in LB media. Briefly, 100 µL of overnight *E. coli* culture was inoculated in 10 mL fresh media and supplemented with 10 mM MgSO_4_ and 10 mM CaCl_2_. The culture was grown until log phase (0.5 OD_600nm_) before infecting it with 100 µL ∼10^8^ PFU/mL T4 phage lysate and incubating overnight at 37°C shaking. The overnight culture was centrifuged 4000 *x g* and the cleared cell lysate treated with 10% (v/v) chloroform to lyse any remaining cells before further centrifugation. The supernatant was passed through a 0.22 µm filter to obtain pure phage lysate.

All phage plaque assays were carried out in *E.coli* DH10B (NEB) using the double layer top agar method. For the binder inhibition assays, DH10B cells were transformed with plasmids carrying an aTc inducible pTET-LbuCas13a^36^ with a co-expressing crRNA targeting T4 soc and an arabinose inducible pBAD vector carrying the binder protein or an empty vector. The strains were grown overnight in LB media containing 100 µg/mL Ampicillin and 35 µg/mL chloramphenicol. A 100 µL aliquot of overnight culture was inoculated into fresh Lennox media and induced at log phase with 2.5 nM aTc and +/- 2% arabinose to induce the expression of the Cas13 effector and binders, respectively. Induced culture was incubated overnight at 37°C shaking. The following day soft Lennox agar (0.5% (w/v) agar) supplemented with 1 mL of overnight culture was poured onto Lennox agar plates supplemented with antibiotics and inducers. Phage titres were calculated by spotting 3 µL drops of serially diluted phage lysate (10^9^-10^1^ PFU/mL) and incubated overnight at 37°C. Plaques were counted after incubation and represented as PFU/mL. Where plaque formation was inhibited, a single plaque was counted on the first dilution with no plaques. Biological triplicates were performed for all binders and the efficiency of plaquing (EOP) in their presence was plotted.

All liquid phage restriction inhibition experiments were performed in Tecan Infinite M200 plate reader. Briefly, the same *E.coli* DH10B strains carrying pTET-LbuCas13a with a co-expressing crRNA and pBAD vector carrying the binder protein were grown overnight at 37°C with appropriate antibiotics. 100 µL of overnight culture was then sub-cultured into fresh Lennox media and induced at log phase with 2.5 nM aTc and +/- 2% arabinose to induce the expression of the Cas effector and binders. Induced culture was incubated overnight at 37°C. Overnight culture was then added at 1:100 dilution to a 96 well plate and monitored on the plate reader until it reached OD 0.1. Wells were then infected with T4 phage at MOI 0.0005 and incubated shaking for 16 hours, collecting OD_600nm_ readings every 20 minutes. Biological triplicates of all assays were performed, and data was plotted using Prism 10. The schematics were created using BioRender.

## Acknowledgements

We would like to thank the Ramaciotti Centre at Monash University, in particular Hariprasad Venugopal, for their support and assistance with data collection on the Talos Arctica. We are also grateful to the Monash Macromolecular Crystallisation Platform (MMCP) for their assistance with crystallization screening. We thank Solve Scientific for their BLI expertise and providing access to a GatorPrime instrument. Special thanks to Marjan Hadian-Jazi for her assistance in evaluating AIcr designs. We acknowledge the MX2 beamline at the Australian Synchrotron (ANSTO) and the Australian Cancer Research Foundation (ACRF) detector. GK was supported by the Snow Medical Research Foundation (SMRF2021-276). Snow Medical were not involved in the design of the study, data collection, analysis, interpretation of the data, the writing, or the decision to submit the article for publication.

## Contributions

C.T., R.G., and G.J.K conceived of and designed the study. C.T. designed, curated, and analysed AIcrs with assistance from R.G. and G.J.K. H.X.C and R.S.B. performed initial AIcr screening with assistance from C.T. H.X.C carried out Cas13 enzyme inhibition assays and analysis with assistance from R.S.B, C.T, B.K.H, R.W.C, and G.J.K. C.T determined and analysed X-ray and cryo-EM structures with assistance from H.X.C, R.S.B. and G.J.K. C.T, H.X.C, and J.D.S performed biophysical experiments with assistance from D.C, F.M, and L.M. J.D.S carried out phage assays with assistance from J.B. All authors contributed to data curation and analysis. C.T. and G.J.K drafted the manuscript. All authors contributed to writing the manuscript.

