## Supplementary Figures and Tables for "*De novo* design of potent CRISPR-Cas13 inhibitors"

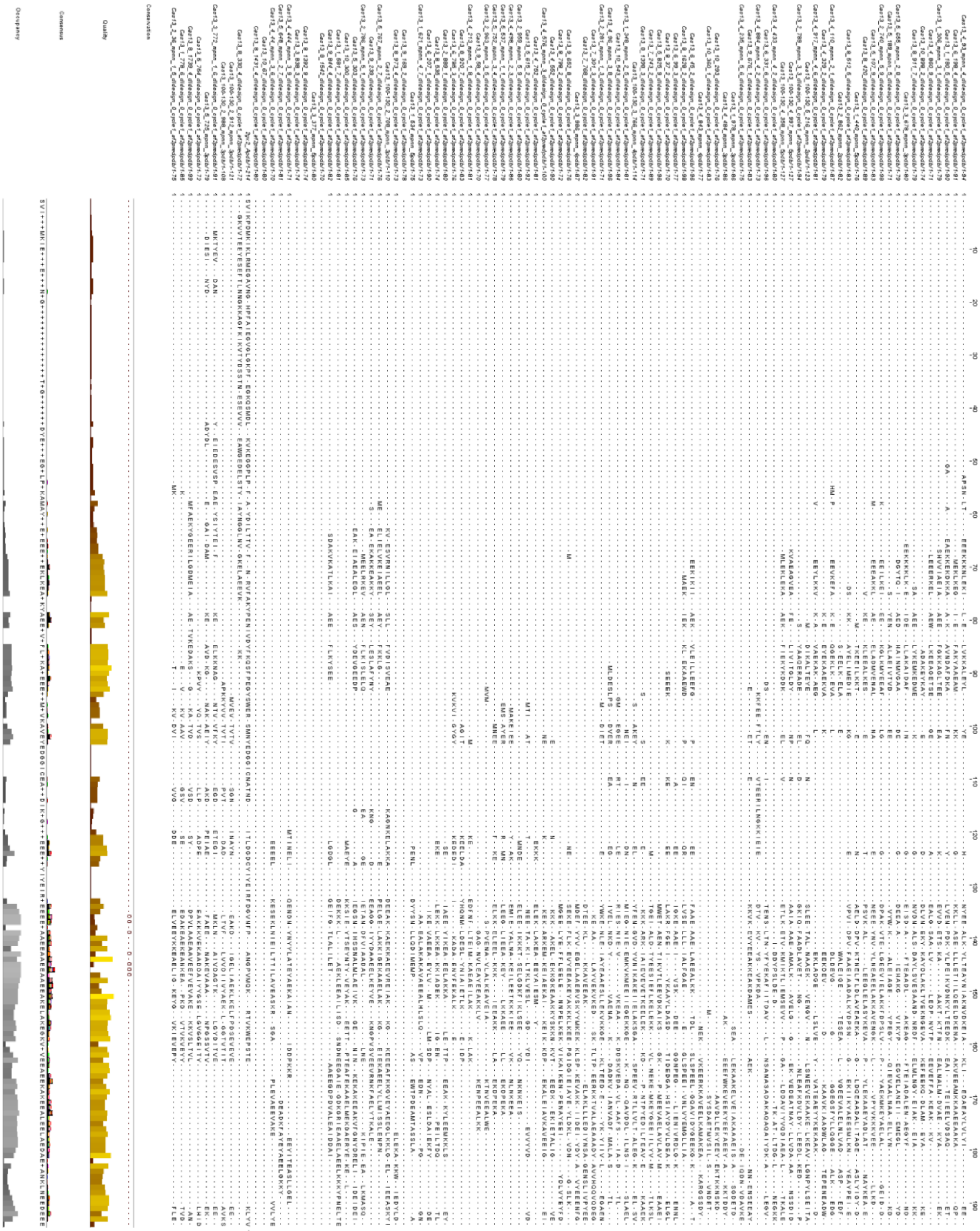

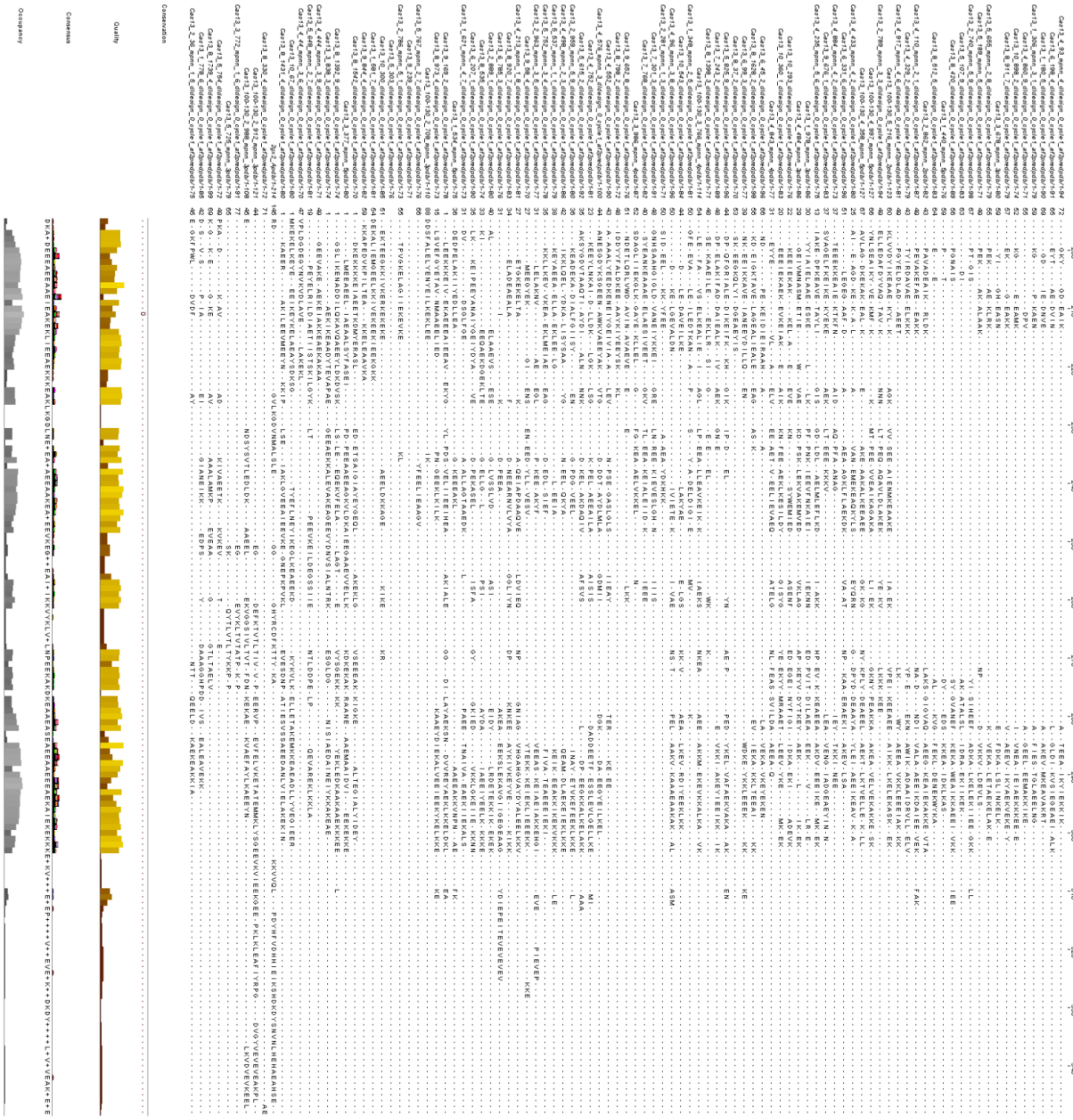

**Supplementary Figure 1. Multiple sequence alignment used for multiple structure alignment (MSTA).** Multiple sequence alignment was generated using FoldMason[1] and visualized using JalView[2], integrating both structural alignment and primary amino acid sequence data from 96 AIcr candidates selected for binding and inhibition of LbuCas13a. The combined structural and sequence-based approach revealed no conserved placement of amino acids in primary sequence and matches the diverse structural landscape.

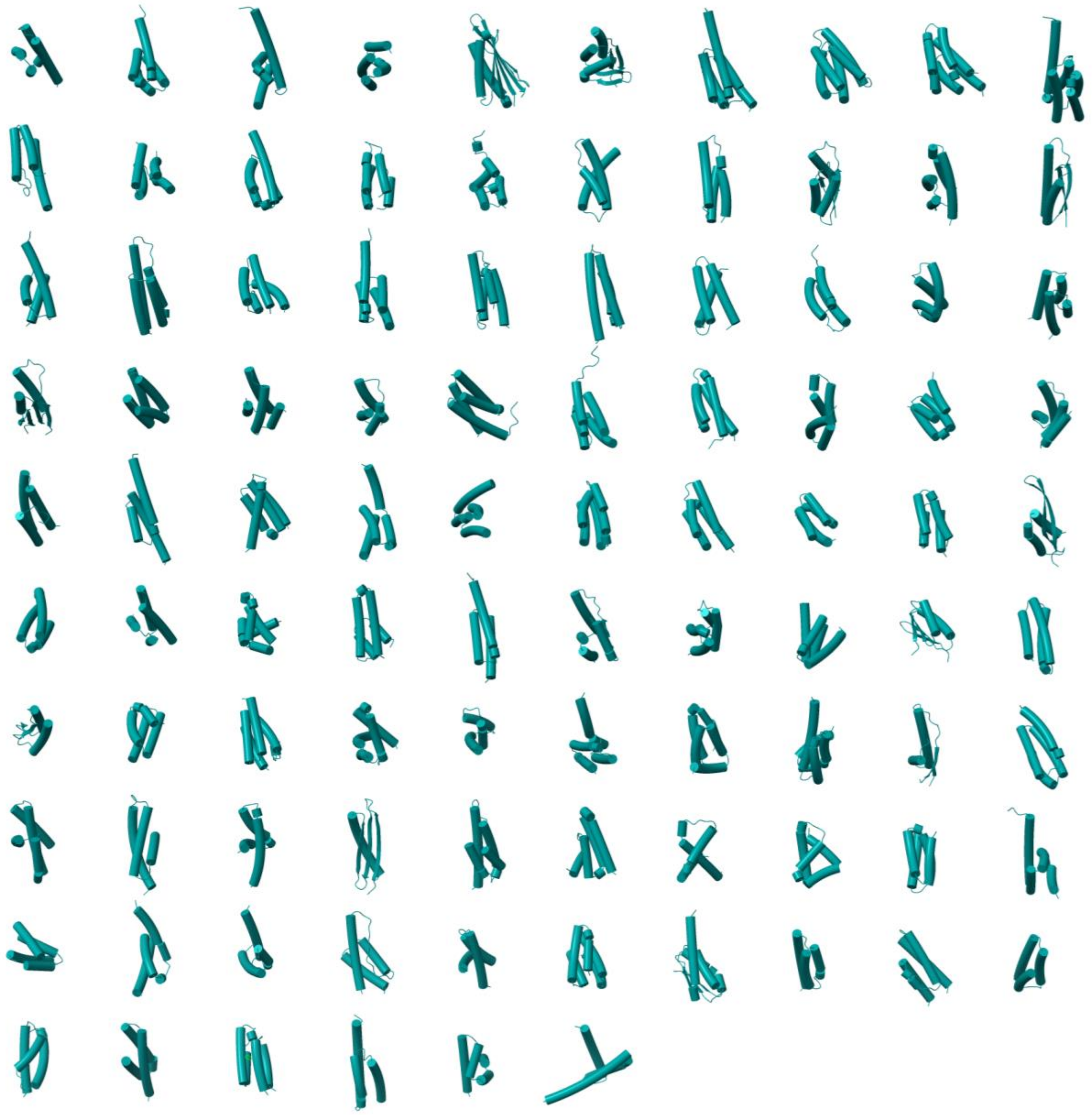

**Supplementary Figure 2. Designed AIcr fold diversity.** AlphaFold2 predictions of the 96 AIcrs tested in this study (cartoon representations shown in teal).

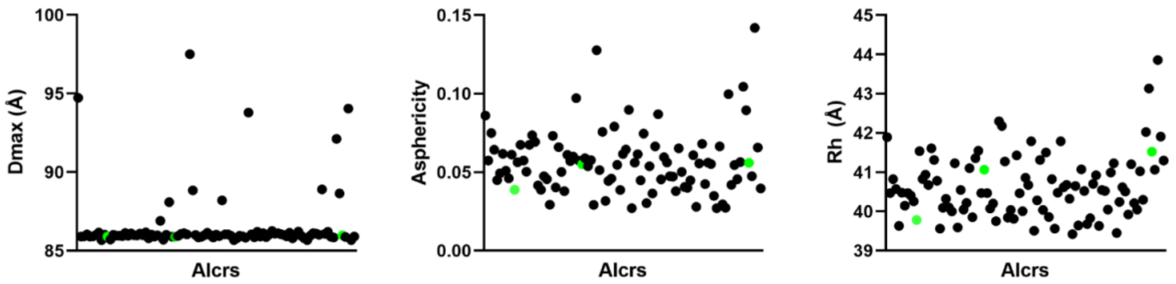


**Supplementary Figure 3. Dimensional and morphological diversity of the 96 AIcrs.** Maximum diameter (Dmax, Å), asphericity, and hydration radius (Rh, Å) were calculated using the MDAnalysis Python package[3]. The majority of Dmax values are approximately 86 Å, asphericity ranges between 0.05 and 0.1, and Rh values are around 40 to 42 Å, indicating that the AI-designed structures exhibit consistent overall size and near-spherical morphology. The green dots refer to AIcrVIA1, AIcrVIA2 and AIcrVIA3.

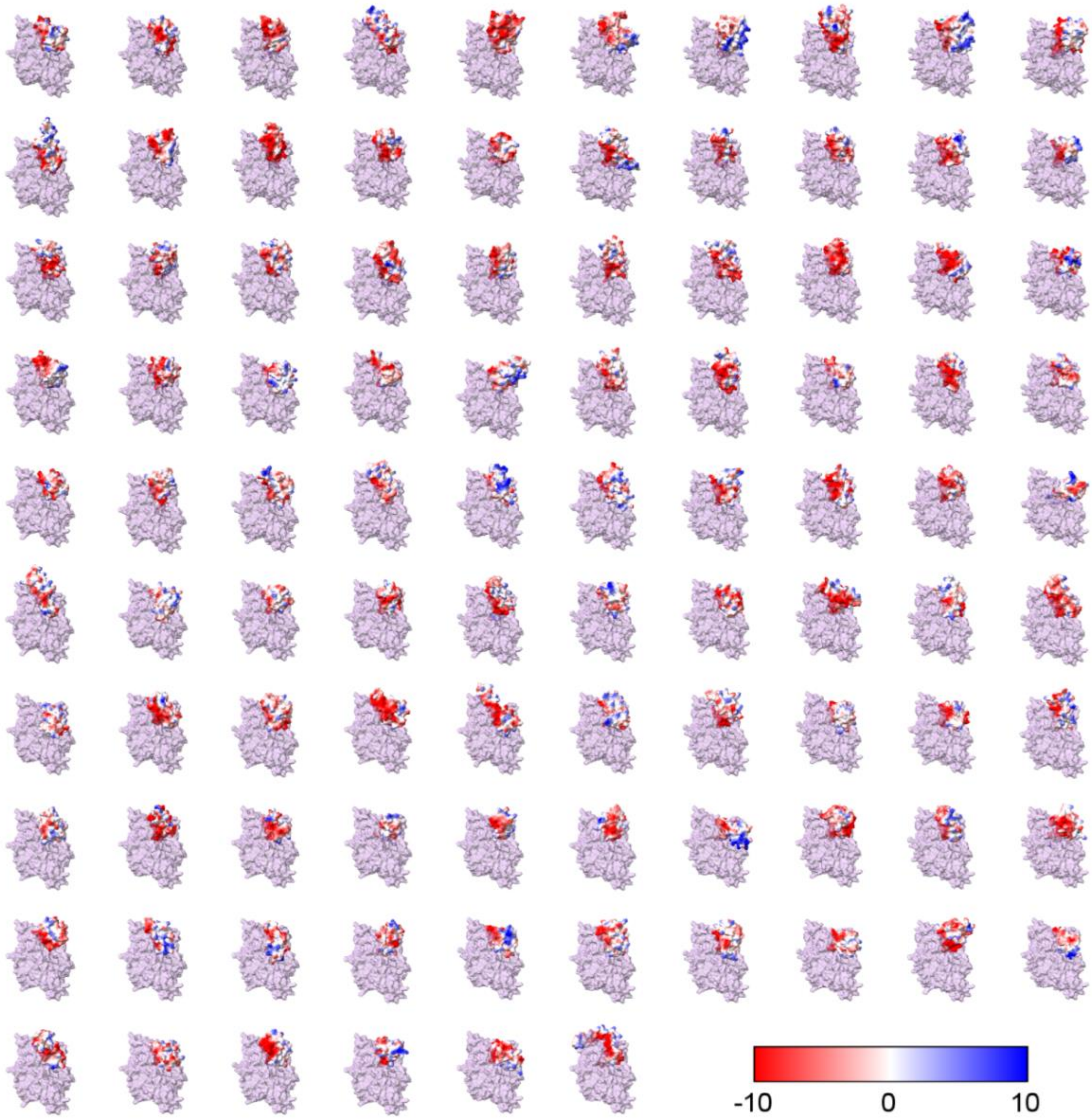


**Supplementary Figure 4. Charge complementarity between AIcr and LbuCas13a active site, mimicking RNA chemical properties**. Coulombic electrostatic surface potential (ChimeraX) of each of the 96 AIcrs (red-blue, surface) in interaction with the HEPN domains of LbuCas13a (surface, purple) used as an input in RF-Diffusion (bottom, transparent). Most designs are predicted to interact via an acidic patch directly with the HEPN domain via charge complementarity.

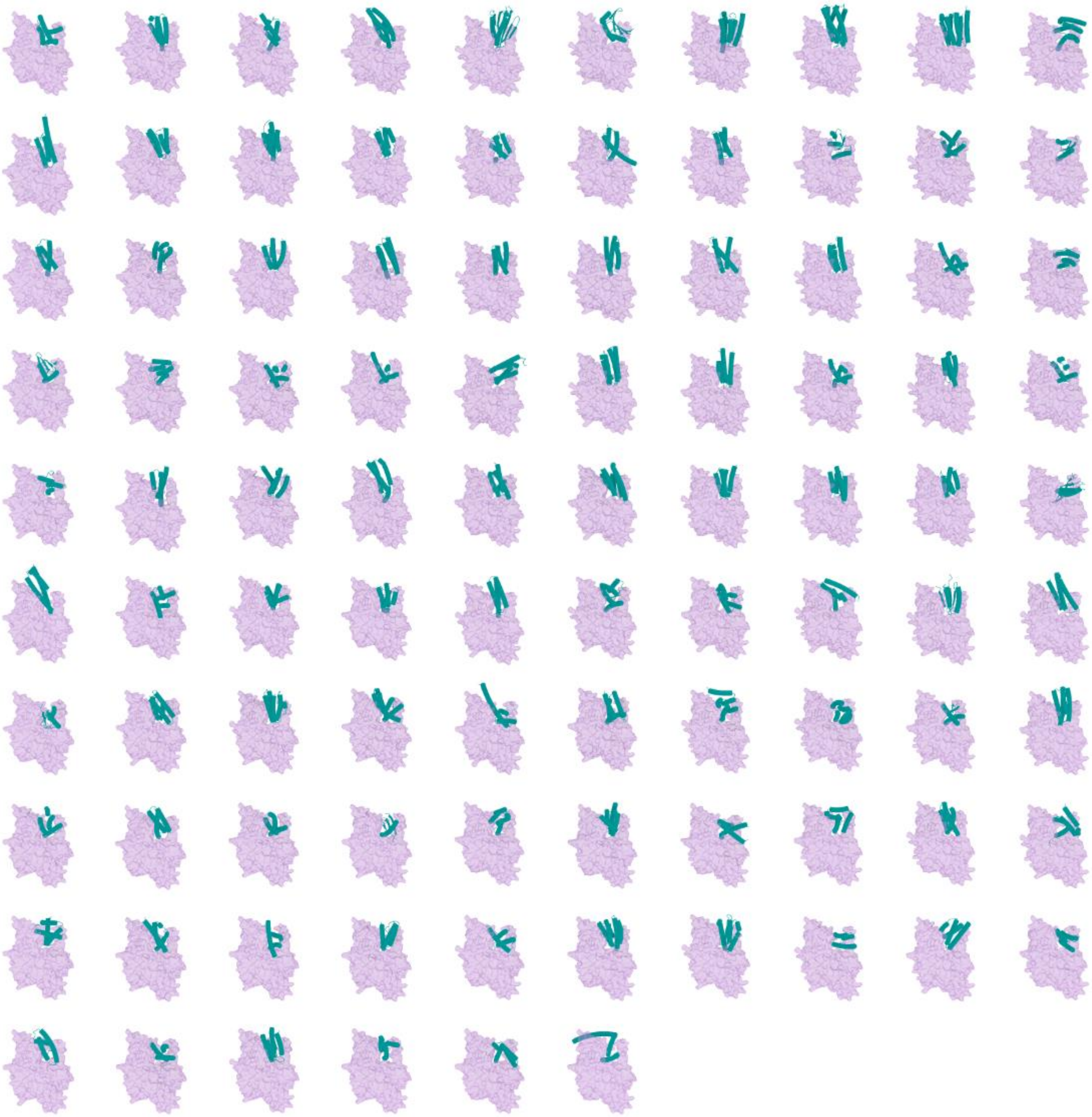

**Supplementary Figure 5. AIcr designs are predicted to interact with the active site of LbuCas13a.** The HEPN domains of LbuCas13a (surface, purple), used as an input in RF-Diffusion, along with each of the 96 different in silico-designed AIcrs (cartoons, teal).

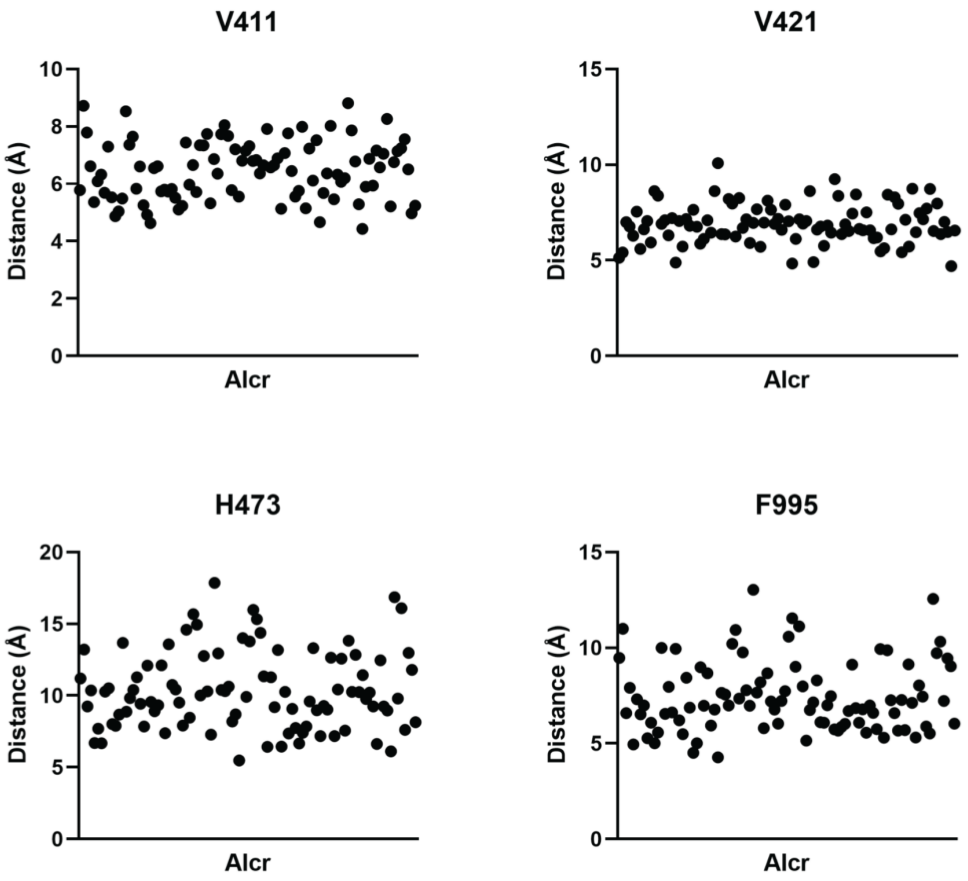


**Supplementary Figure 6. Distance between key hotspot residues of LbuCas13a and the 96 potential AIcr designs.** Minimum distances between the key hotspot residues used for the AIcrs design (V411, V421, H473, and F995) in the LbuCas13a active site and each of the 96 AIcr designs. Most AIcrs are predicted to be between 5 – 15 Å, reflecting potential interactions at the LbuCas13a active site.

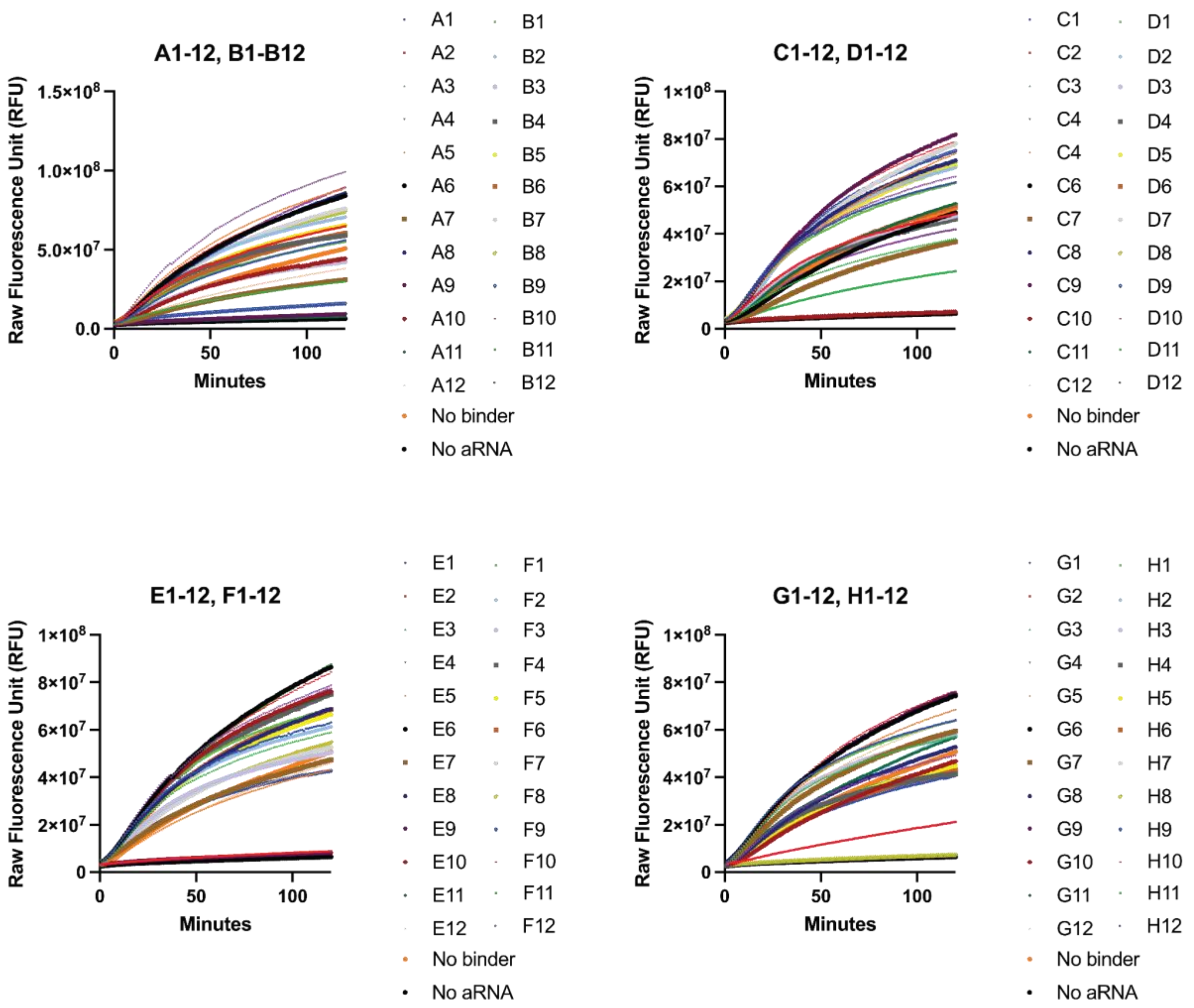


**Supplementary Figure 7. LbuCas13a HEPN nuclease activity assay in the presence of cell-free expressed AIcrs.** Each AIcr (A1-H12 in a 96 well plate) was assayed with LbuCas13a-crRNA-aRNA in the presence of labelled reporter RNA. Activity from LbuCas13a leads to increased fluorescence (RFU) over time (mins).

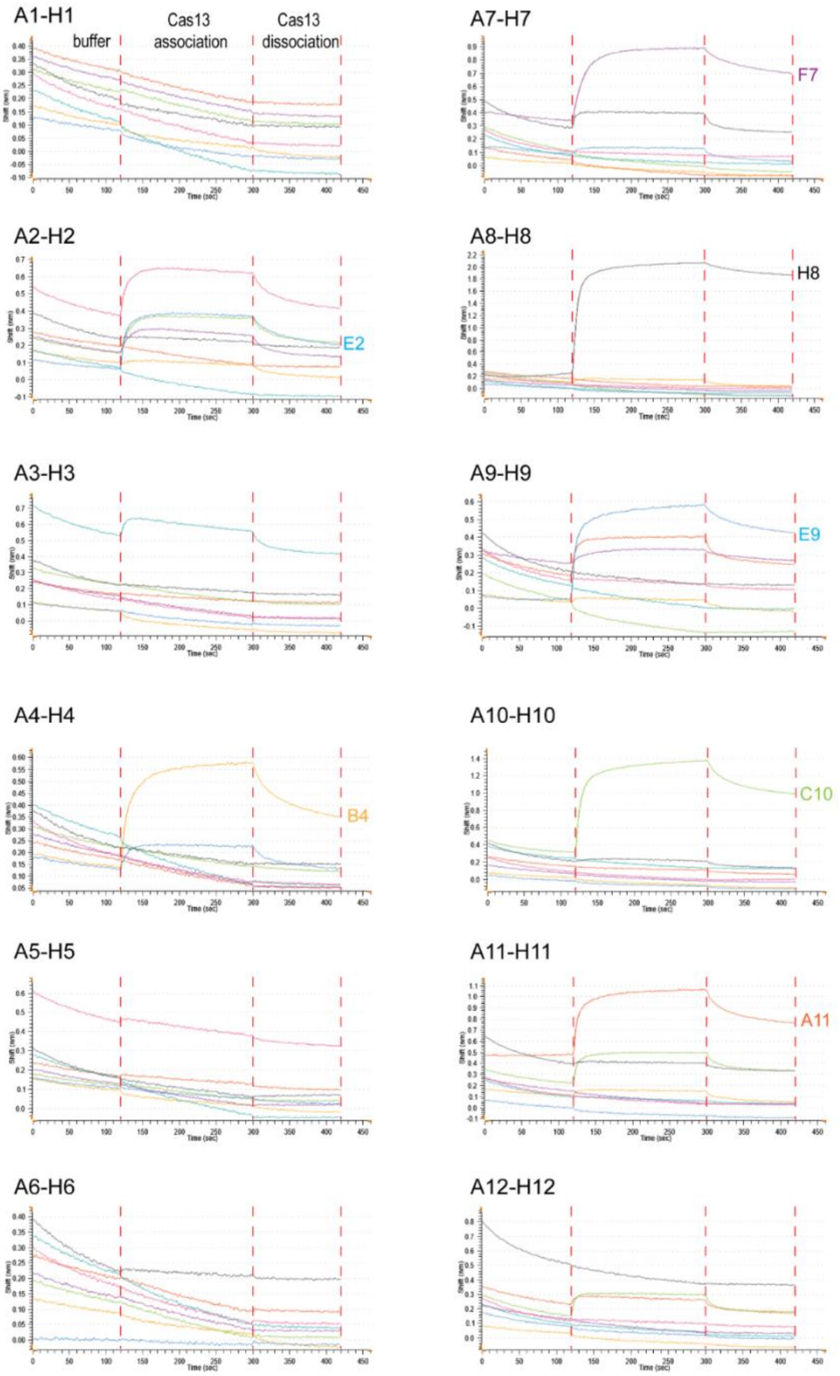


**Supplementary Figure 8. Raw biolayer interferometry data for AIcr binding to the LbuCas13a-crRNA-aRNA complex.** Each experimental panel shows a plate column of eight AIcr designs (A1-H1, A2-H2, etc.) loaded onto Anti-His probes, followed by washing with buffer Q. The association phase is represented by binding of LbuCas13a-crRNA-aRNA, and the subsequent dissociation phase is shown after washing with buffer Q. The raw data were normalized by subtracting the signal from a ‘no AIcr bound’ control experiment using the same probes, and the difference is depicted for each phase (wash, association, and dissociation).

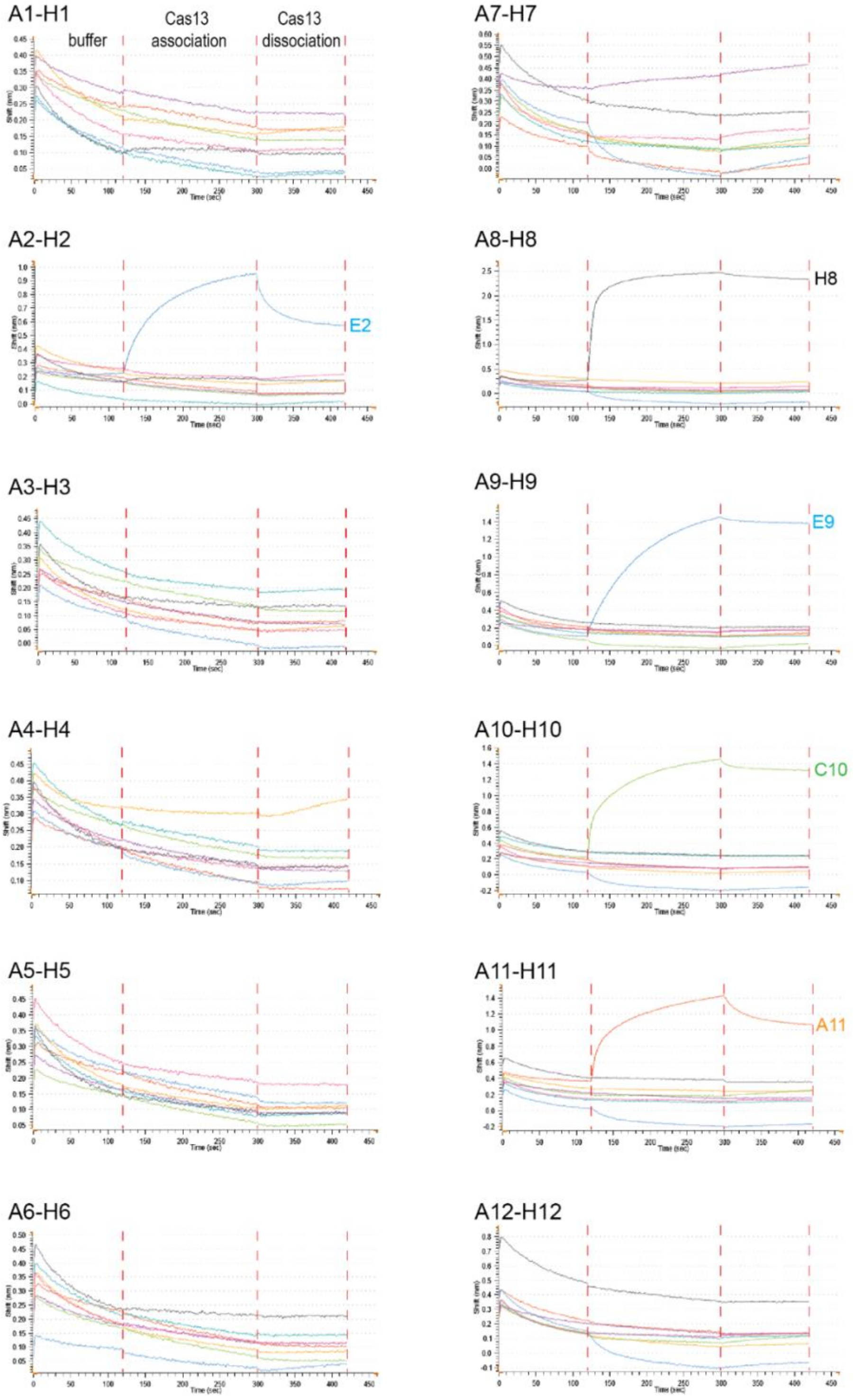


**Supplementary Figure 9. Raw biolayer interferometry data for AIcr binding to the LbuCas13a-crRNA complex.** Each panel represents data from a plate column containing eight AIcr designs (A1-H1, A2-H2, etc.) loaded onto Anti-His probes, followed by a wash with buffer Q. The association phase shows the binding of LbuCas13a-crRNA to AIcr, while the dissociation phase follows after an additional wash with buffer Q. The signals were normalized by subtracting the data from a ‘no AIcr bound’ control experiment using the same probes. The difference in signal is shown for the wash, association, and dissociation phases.

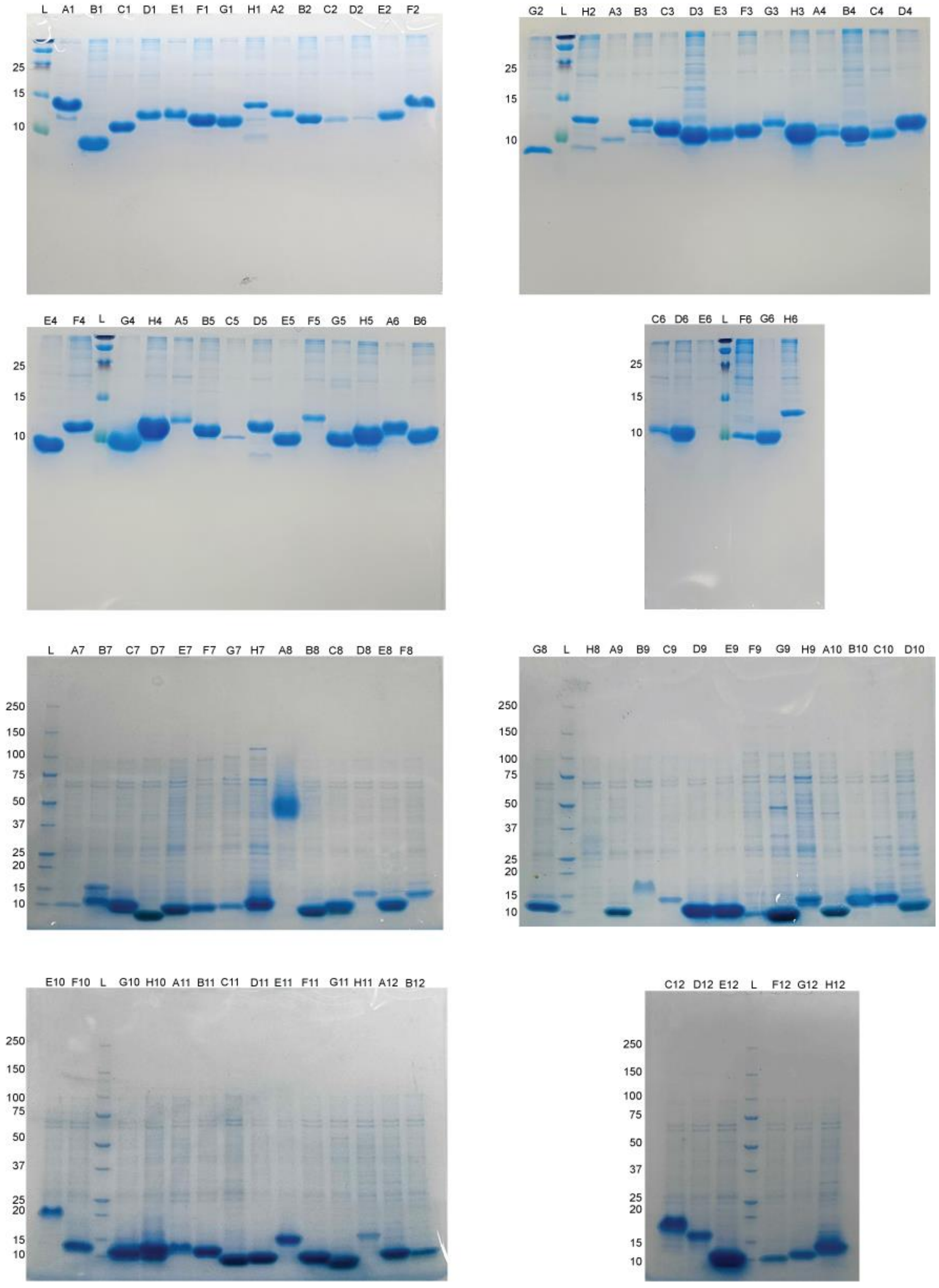


**Supplementary Figure 10. Small-scale purification of AIcrs.** AIcrs were expressed and purified in small scale followed by batch-based affinity chromatography and SDS-PAGE. (TOP) 12% (w/v) SDS-PAGE gels, (BOTTOM) 4-20% (w/v) gradient SDS-PAGE gels. The ladder (L) molecular weights (MW) are in kDa (left side of gel).

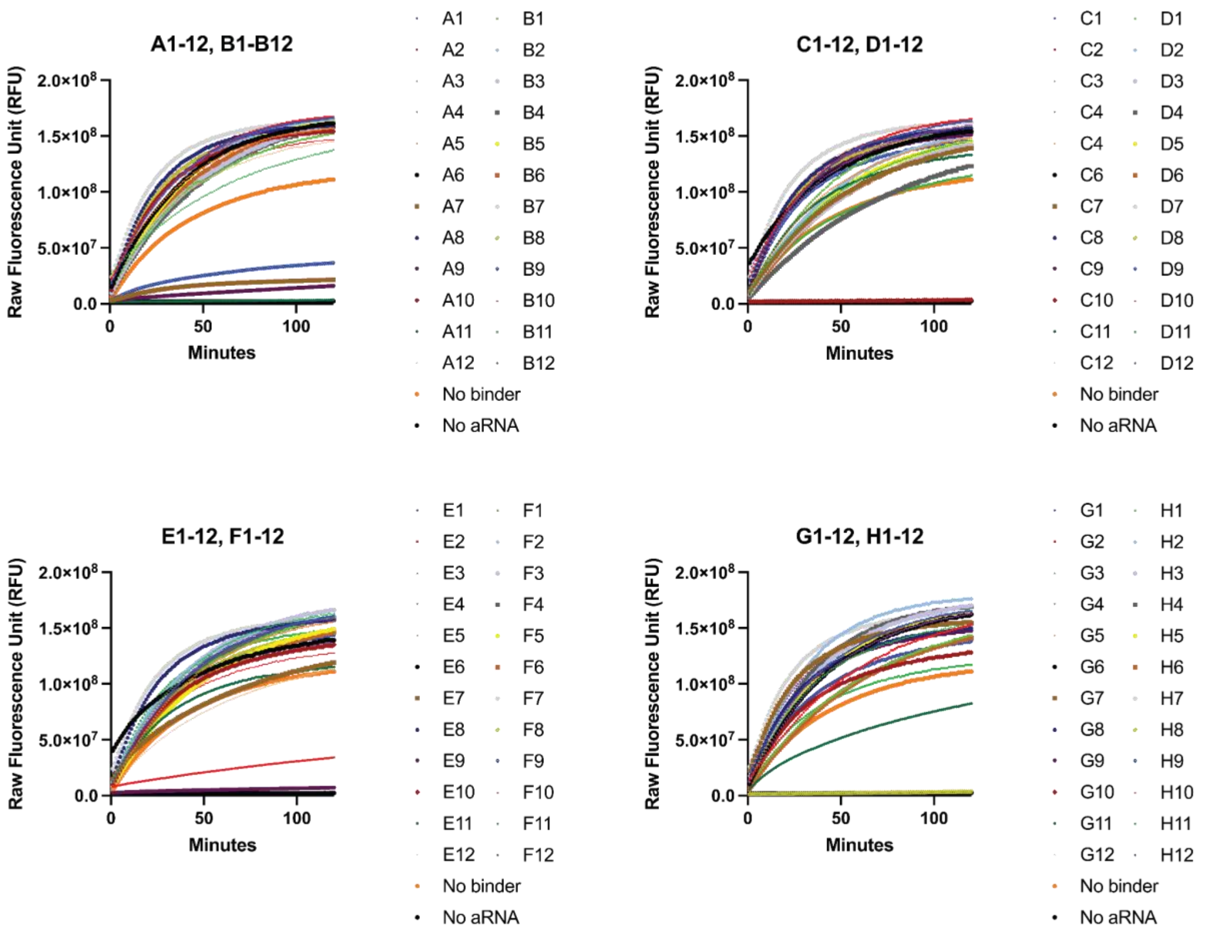


**Supplementary Figure 11. LbuCas13a HEPN nuclease activity assay in the presence of semi-purified AIcrs.** Each AIcr (A1-H12 in a 96 well plate) was assayed with LbuCas13a-crRNA-aRNA in the presence of labelled reporter RNA. Activity from LbuCas13a leads to increased fluorescence (RFU) over time (mins).

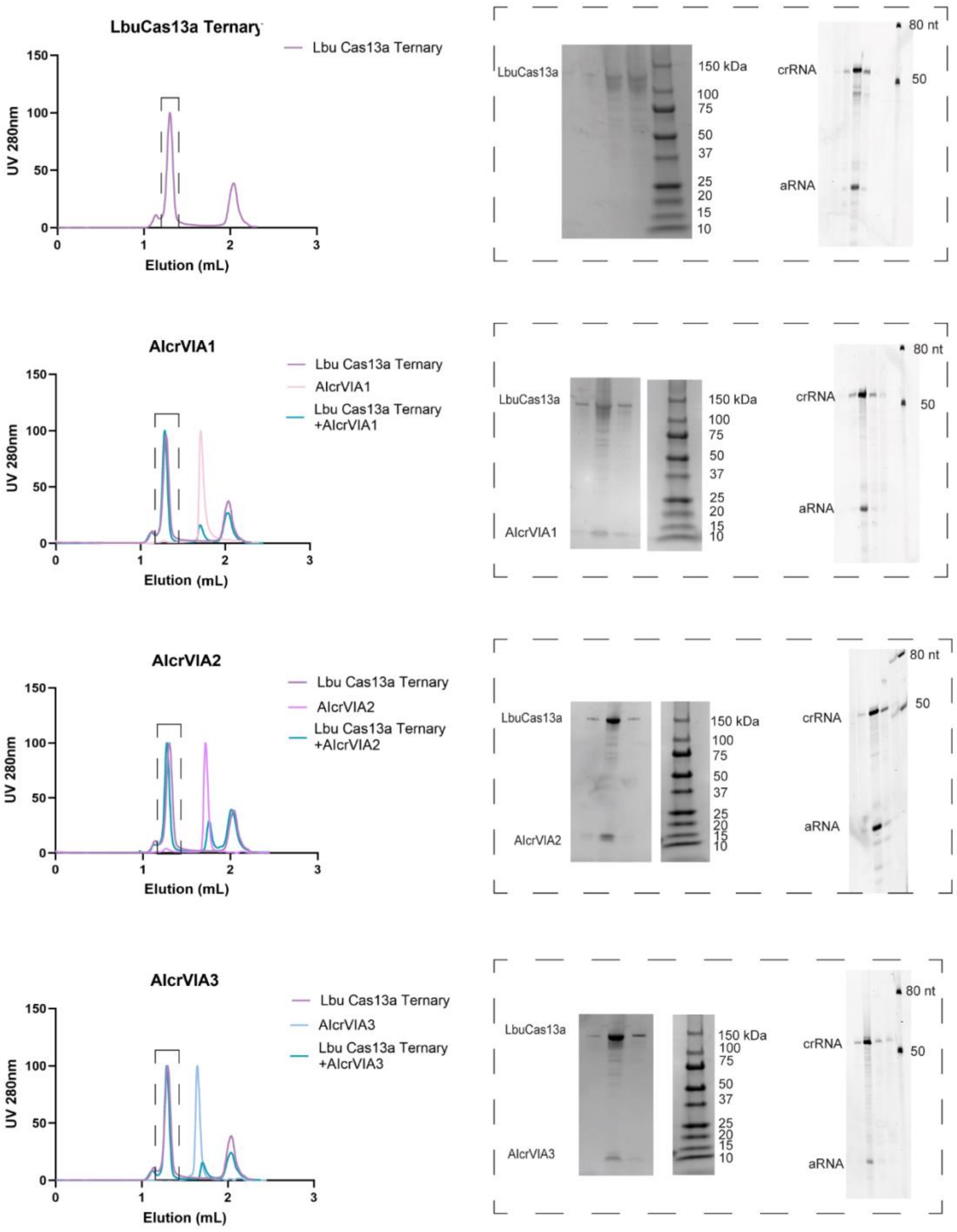


**Supplementary Figure 12. AIcrVIA1, AIcrVIA 2, and AIcrVIA3 interact with LbuCas13a.** Gel filtration trace (left) and corresponding SDS-PAGE and urea gels (right) are shown for protein and RNA detection throughout the elution process. The red dashed rectangles on the gel filtration chromatogram indicate the fractions corresponding to the SDS-PAGE results, while the black dashed rectangles highlight the fractions analyzed by urea gel electrophoresis. Coelution of LbuCas13a with the AIcr binders indicates an interaction between the AIcrVIA designs (A1, A2, and A3) and LbuCas13a.

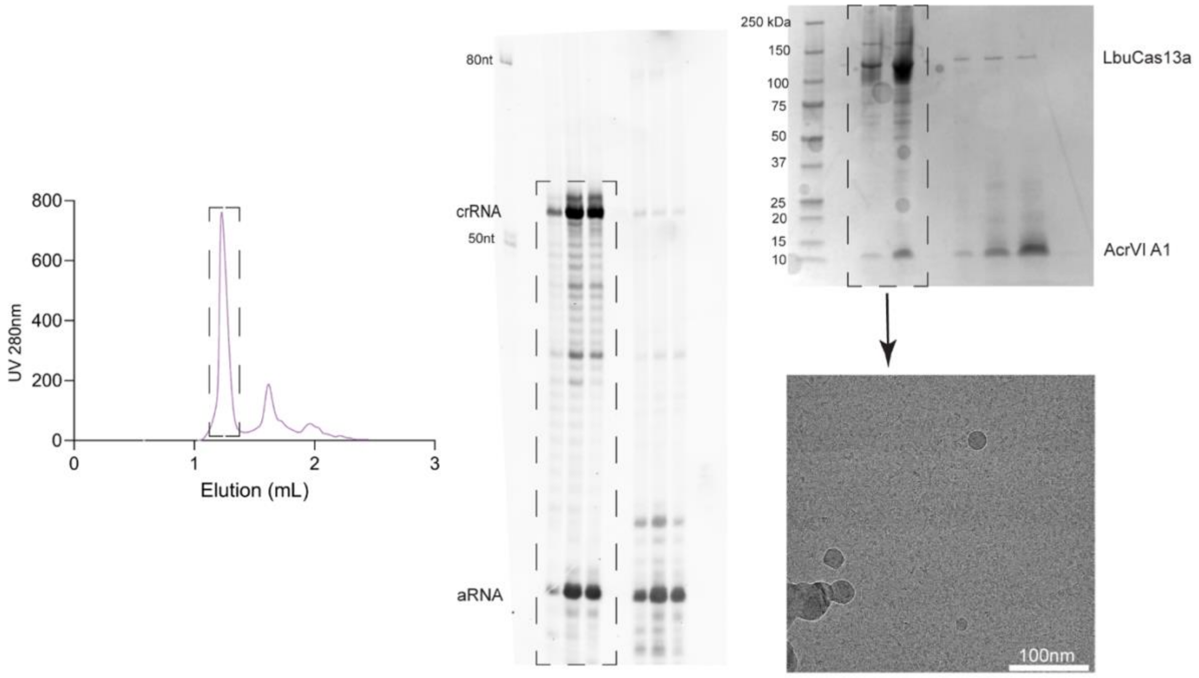


**Supplementary Figure 13. LbuCas13a-AIcrVIA1 complex preparation for cryo-EM.** The left panel shows the gel filtration profile of the ternary LbuCas13a-AIcrVIA1 complex. Protein, crRNA, and aRNA components are verified by SDS-PAGE (top right) and urea-PAGE (middle right), confirming co-elution of the components for optimal cryo-EM sample preparation (black dashed rectangles). The bottom right panel displays a cryo-EM micrograph.

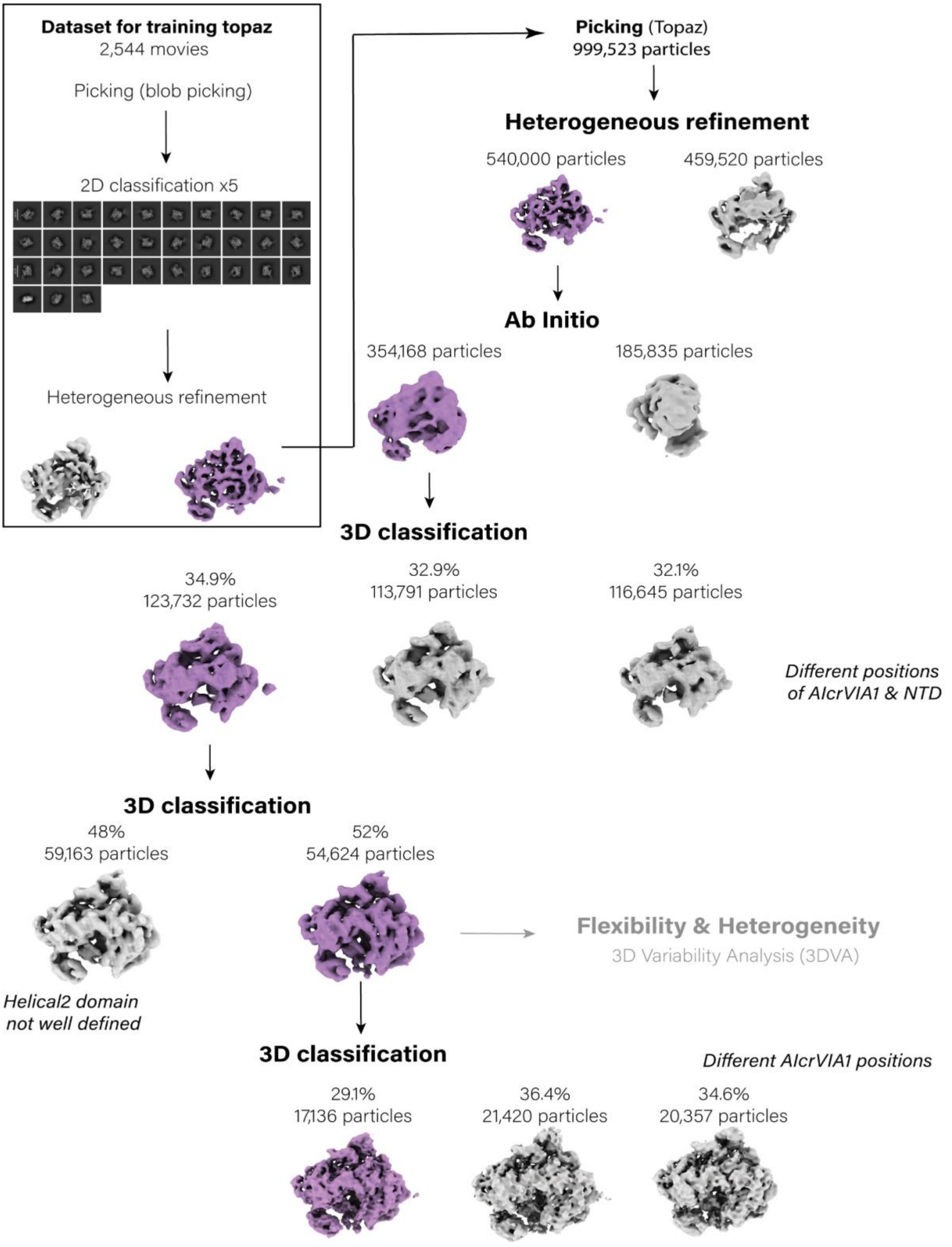


**Supplementary Figure 14. Step-by-Step Cryo-EM Workflow for data processing.** A detailed schematic of the cryo-electron microscopy (cryo-EM) data processing pipeline for LbuCas13a-crRNA-aRNA in complex with AlcrVIA1, from sample preparation through to image acquisition and analysis.

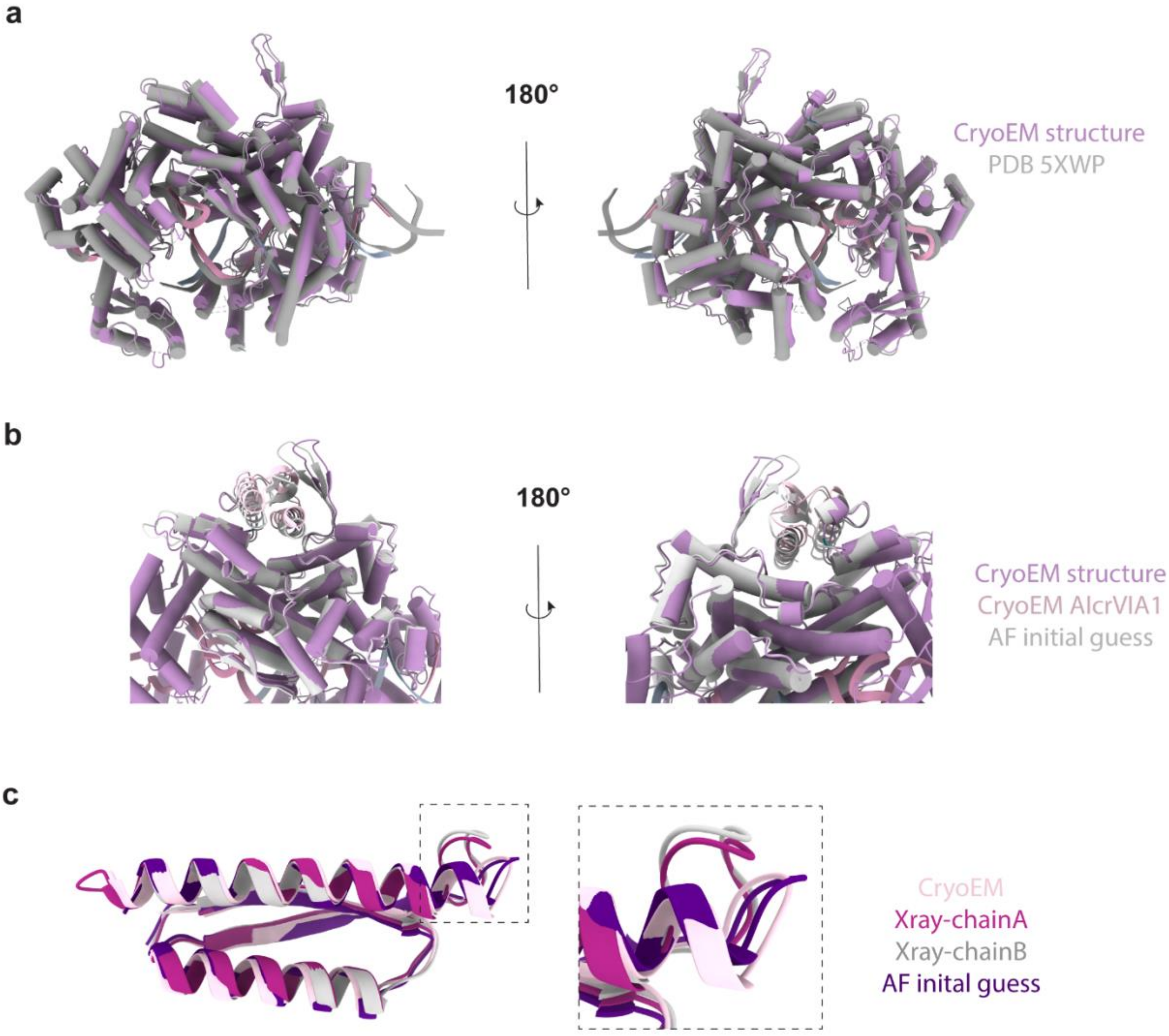


**Supplementary Figure 15. Cryo-EM atomic model of the LbuCas13a ternary complex bound to AlcrVIA1 resembles the RF-diffusion prediction. a.** Superposition of the LbuCas13a ternary complex atomic model obtained through cryo-EM (cartoon, purple) and the X-ray crystallography structure of the same complex (PDB: 5XWP, cartoon, grey). The structures are highly similar, highlighting the overall structural conservation of the Cas13 ternary complex bound to AlcrVIA1. **b.** Comparison of the cryo-EM model (purple LbuCas13a, pink AlcrVIA1) with the AI-predicted model. The models show striking similarity, with differences in the β-turn region (residues 401-421) and a slight rotation in the AlcrVIA1 position. These deviations can be attributed to the inherent flexibility observed in our heterogeneity studies (**Supplementary Movie 1**). **c.** Cartoon representation of AlcrVIA1 from the cryo-EM atomic model (light pink), X-ray structures (chain A, pink; chain B, grey), and AF initial guess results (violet). The overall alignment between the four structures is consistent, except for a loop region (indicated by black dashed lines), where the cryo-EM structure resembles the AF initial guess more closely than the X-ray structures. This difference may be due to constraints imposed by crystal packing.

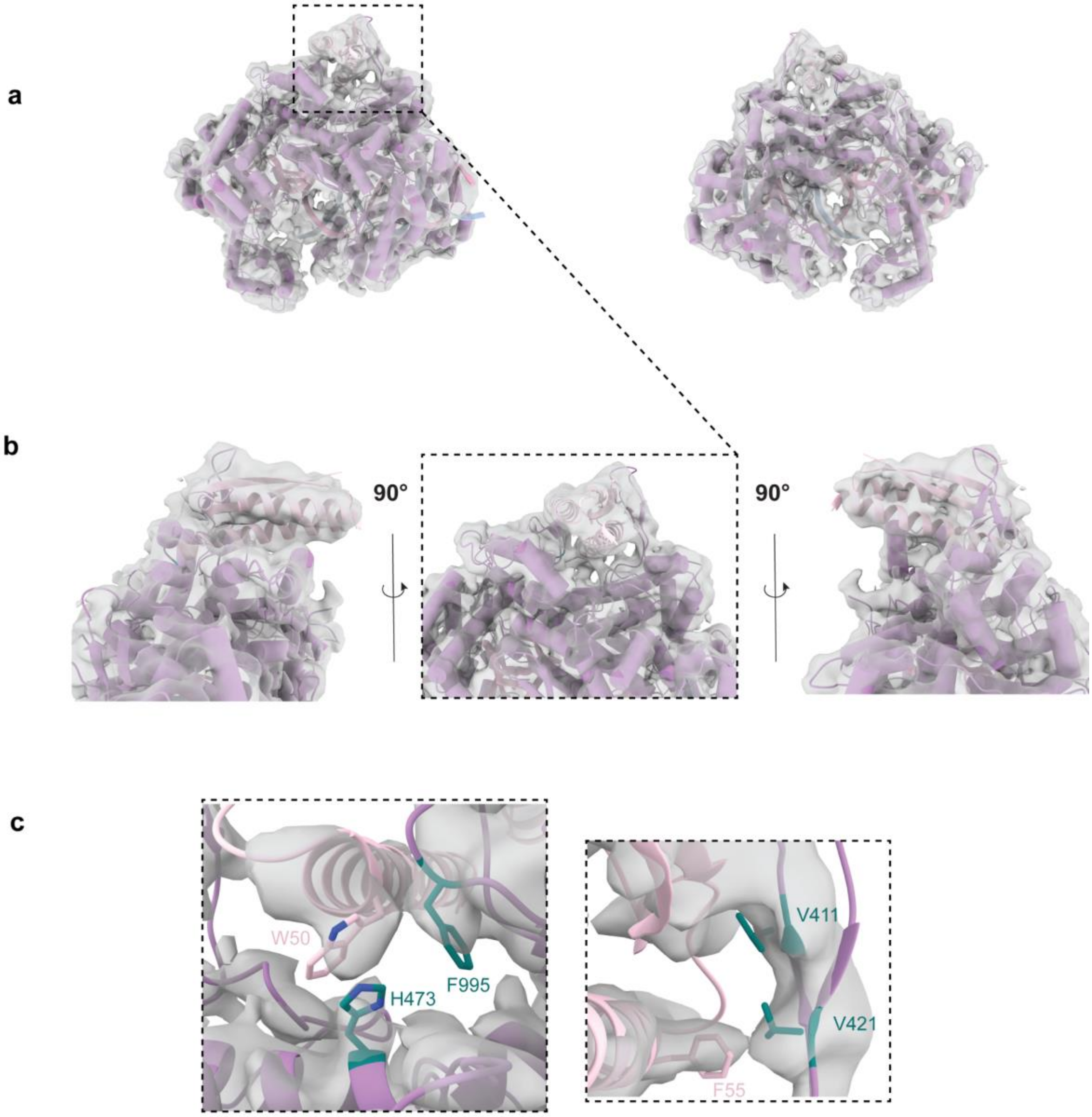


**Supplementary Figure 16. Mechanism of AIcrVIA1 inhibition aligns with design predictions. a.** Overall fit of the cryo-EM map (transparent surface) for the LbuCas13a-crRNA-aRNA complex bound to AIcrVIA1 (indicated by black dashed lines) atomic model. **b.** Zoomed view of the HEPN1 and HEPN2 domains of LbuCas13a (purple cartoon). AIcrVIA1 is positioned consistently with AI design predictions. **c.** Stick representation of the key interaction hotspots targeted during the AI design workflow (teal). These hotspots are located near aromatic residues, highlighting strong interactions between LbuCas13a and AIcrVIA1. However, structural heterogeneity in this region of the cryo-EM density map (transparent surface) limits high-resolution features.

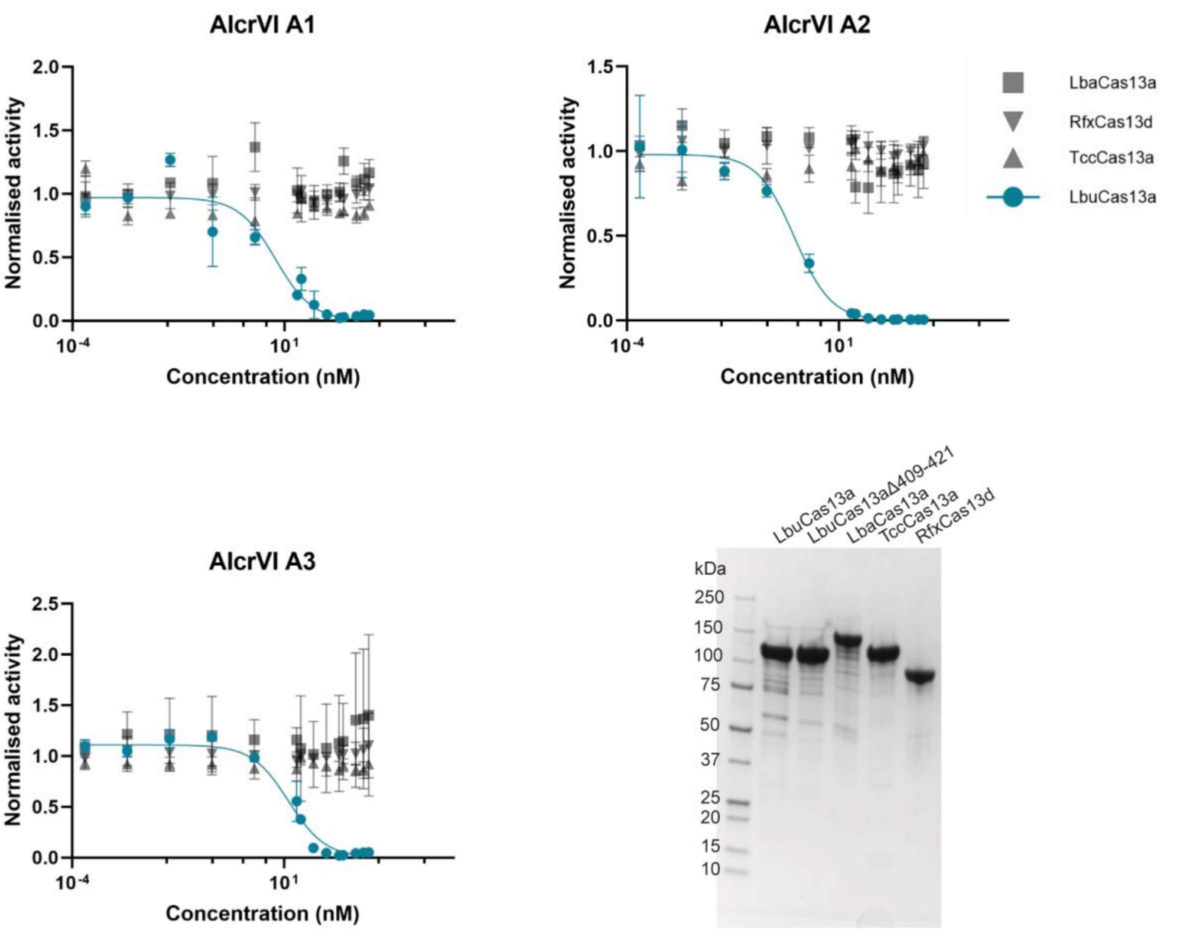


**Supplementary Figure 17. AIcrVIA1, AIcrVIA2, and AIcrVIA3 are specific to LbuCas13a.** Activity assays were conducted for LbuCas13a, LbaCas13a, TccCas13a, and RfxCas13d in the presence of AIcrVIA1 (top left), AIcrVIA2 (top right), or AIcrVIA3 (bottom left). While the nuclease activity of LbuCas13a is nearly abolished by these AIcrVIA inhibitors, the activity of the other Cas13 proteins remains unaffected, demonstrating the specificity of inhibition. The purity of all Cas13 proteins was confirmed by SDS-PAGE analysis (bottom right).

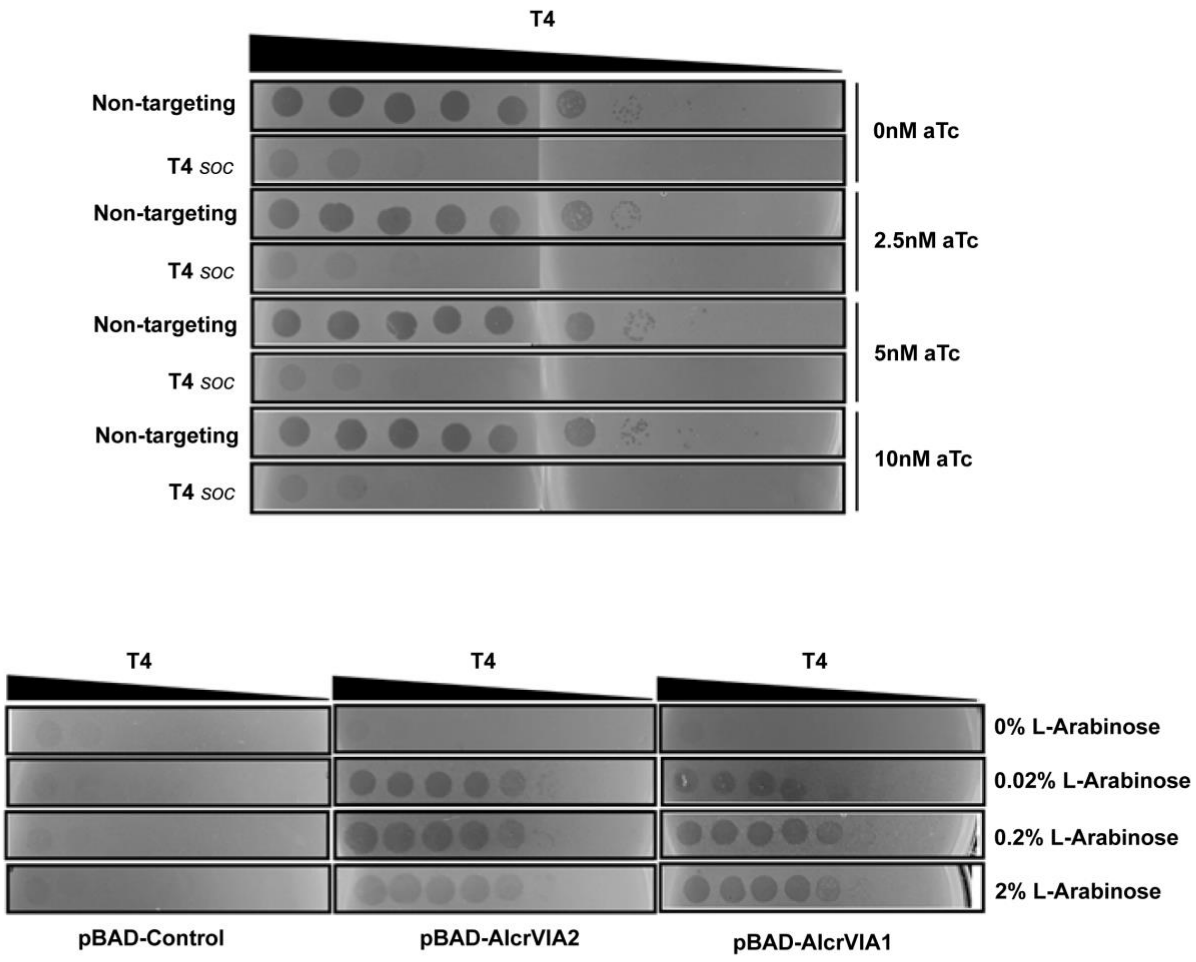


**Supplementary Figure 18. Optimization of protein expression conditions for the phage assay. a.** Plaque assay optimization of LbuCas13a expression and crRNA production in E. coli using aTc induction. This assay was performed with a plasmid carrying an aTc-inducible LbuCas13a vector and either a constitutively expressed non-targeting (NT) crRNA or a T4 soc-targeting crRNA. No toxicity was observed upon induction. (***NT****: non-targeting crRNA;* ***T4 soc:*** *T4 soc-targeting crRNA*.) **b.** Optimization of AIcrVIA1 and AIcrVIA2 expression with increasing arabinose concentrations. Results indicated that 2% arabinose provided optimal expression levels. Optimization of AIcrVIA3 was not performed, as 2% arabinose was deemed sufficient based on these results.

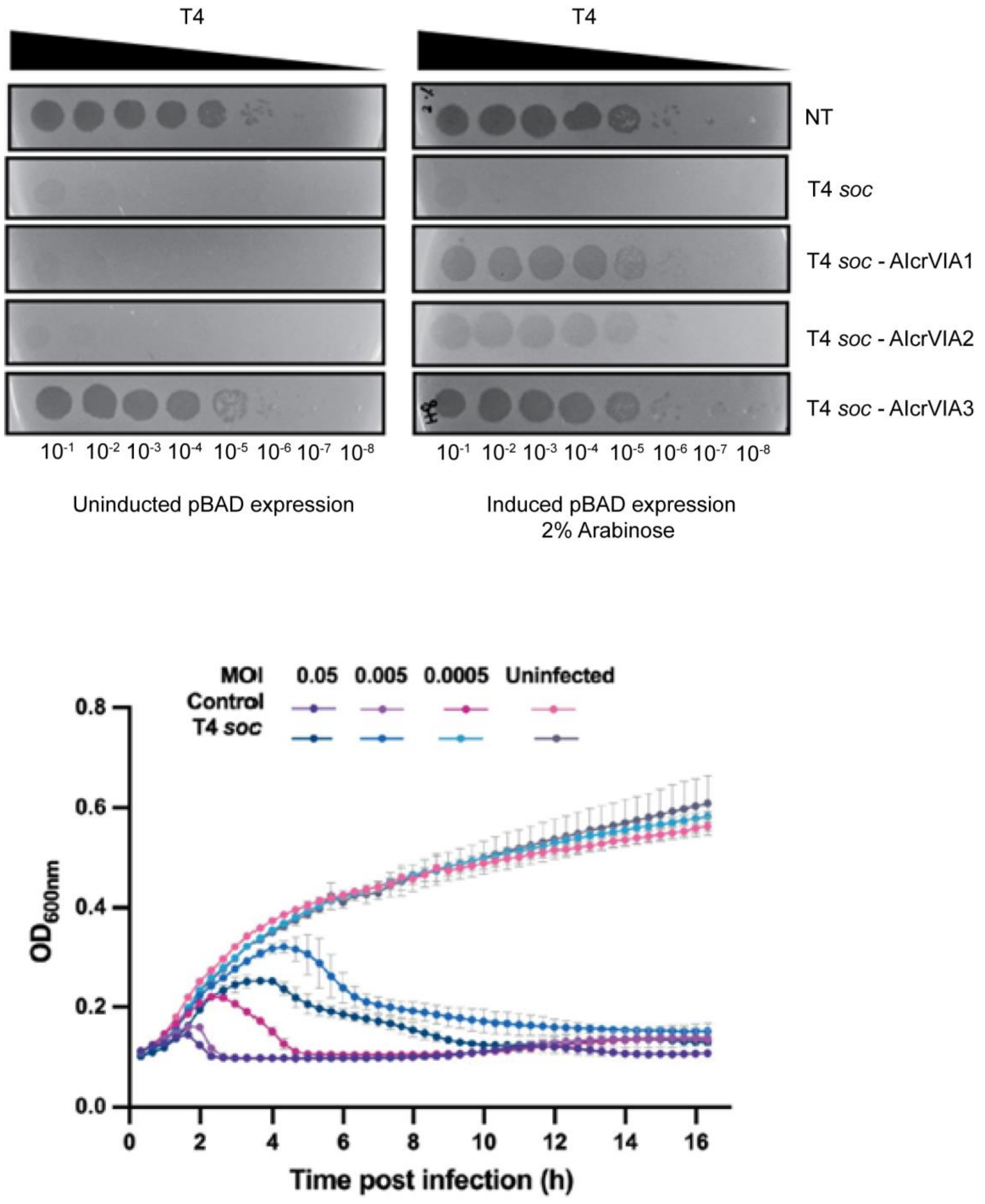


**Supplementary Figure 19. Evaluation of AIcr-mediated inhibition and determination of optimal MOI for phage resistance. a.** Comparison of LbuCas13a inhibition by AIcrs under induced (arabinose) and uninduced conditions using plaque assay. Arabinose induction effectively suppresses LbuCas13a activity, demonstrating active inhibition by AIcrs. **b.** Determination of the multiplicity of infection (MOI) required for LbuCas13a to confer resistance against T4 phage infection in liquid culture. The optimal MOI was identified by monitoring infection dynamics and escape efficiency.

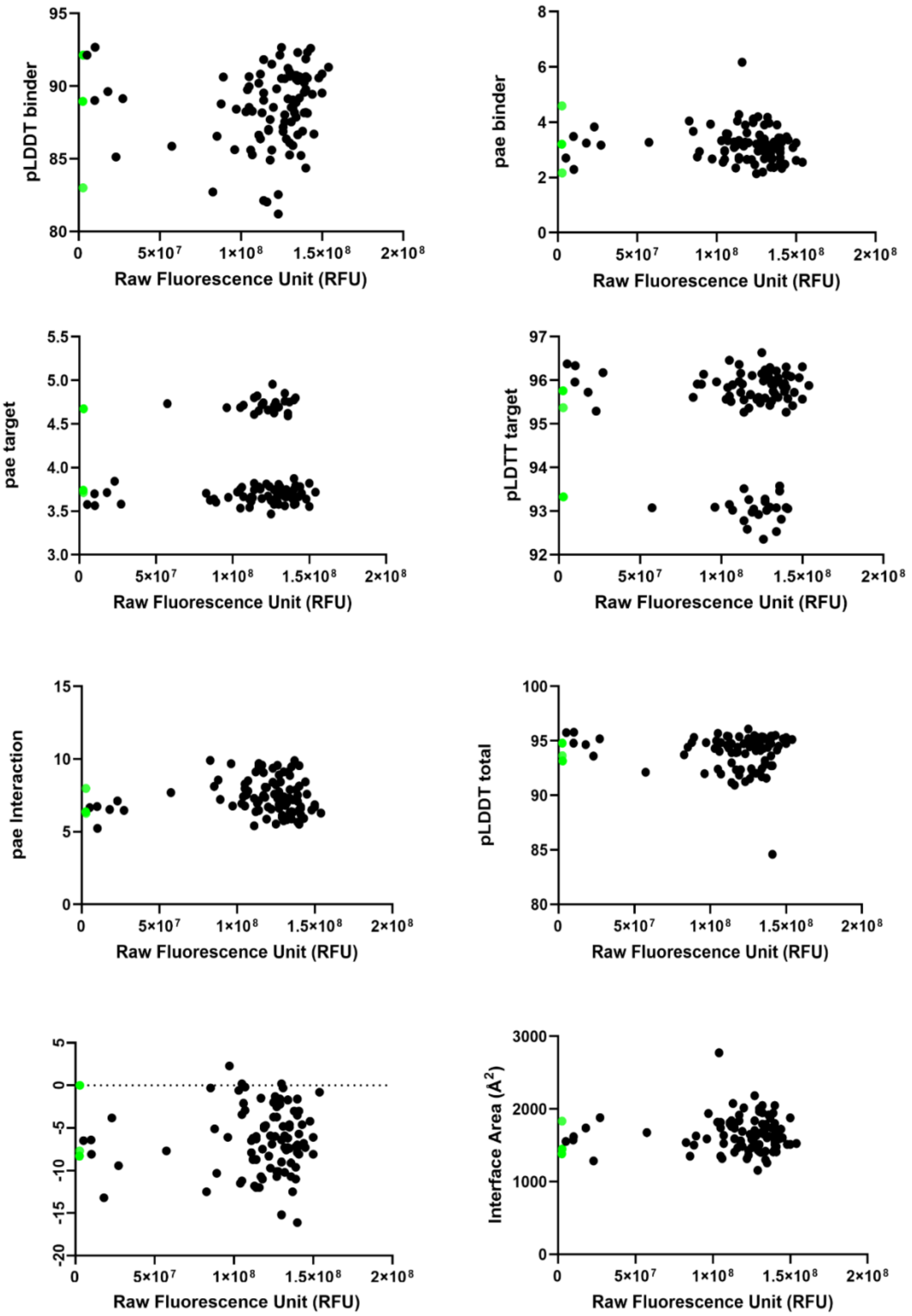


**Supplementary Figure 20: Comparative analysis of PAE, pLDDT scores, predicted ΔG, and interaction surface area.** Correlation analysis of these parameters with LbuCas13a-crRNA-aRNA activity in the presence of 96 AIcrVIA variants. Data points representing AIcrVIA1, VIA2, and VIA3 are highlighted in purple. No correlation was observed between the inhibition capacity of AIcrVIA variants and any of these parameters (PAE, pLDDT scores from AF2 initial guess; predicted ΔG and interaction surface area calculated using PISA).

| | |
|---|---|
| Wavelength | 0.95365 |
| Resolution range | 42.46 - 1.938 (2.007 - 1.937) |
| Space group | P 21 21 21 |
| Unit cell | 33.403 62.9 84.927 90 90 90 |
| Total reflections | 175398 (16269) |
| Unique reflections | 13829 (1274) |
| Multiplicity | 12.7 (12.8) |
| Completeness (%) | 99.45 (95.28) |
| Mean I/sigma(I) | 16.22 (1.99) |
| Wilson B-factor | 36.74 |
| R-merge | 0.09105 (0.9414) |
| R-meas | 0.0951 (0.9805) |
| R-pim | 0.02696 (0.2698) |
| CC1/2 | 0.999 (0.812) |
| CC* | 1 (0.947) |
| Reflections used in refinement | 13820 (1273) |
| Reflections used for R-free | 1383 (127) |
| R-work | 0.1992 (0.2923) |
| R-free | 0.2461 (0.3481) |
| CC(work) | 0.962 (0.841) |
| CC(free) | 0.929 (0.777) |
| Number of non-hydrogen atoms | 1341 |
| macromolecules | 1258 |
| ligands | 1 |
| solvent | 82 |
| Protein residues | 154 |
| RMS(bonds) | 0.003 |
| RMS(angles) | 0.57 |
| Ramachandran favored (%) | 98.67 |
| Ramachandran allowed (%) | 1.33 |
| Ramachandran outliers (%) | 0 |
| Rotamer outliers (%) | 0 |
| Clashscore | 2.8 |
| Average B-factor | 43.25 |
| macromolecules | 43.13 |
| ligands | 69.61 |
| solvent | 44.73 |
| Number of TLS groups | |

**Supplementary Table 1. X-ray Structure Validation and Refinement Statistics Using PHENIX**

| **Data collection and Processing** | | |
|---|---|---|
| Magnification | 150K | |
| Voltage | 200kV | |
| Electron Exposure | 50 $e/Å^2$ | |
| Defocus range | -0.8 -1.4 |  |
| Pixel size | 0.9359 | |
| Initial particles images | 999,523 | |
| Final particle images | 20,357 | |
| **Model** | | |
| **Bonds (RMSD)** | | |
| Length (Å) (# > 4s) | 0.005 (0) | |
| Angles (°) (# > 4s) | 0.728 (2) | |
| MolProbity score | 2.37 | |
| Clash score | 19.04 | |
| **Ramachandran plot (%)** | | |
| Outliers | 0 | |
| Allowed | 2.39 | |
| Favored | 97.61 | |
| **Rama-Z (Ramachandran plot Z-score, RMSD)** | | |
| whole (N = 1171) | 0.49 (0.24) | |
| helix (N = 776) | 1.00 (0.18) | |
| sheet (N = 72) | -0.12 (0.61) | |
| loop (N = 323) | -1.11 (0.33) | |
| Rotamer outliers (%) | 4.69 | |
| Cß outliers (%) | 0.09 | |
| **ADP (B-factors)** | | |
| Iso/Aniso (#) | 11352/0 | |
| **min/max/mean** | | |
| Protein | 89.20/256.04/161.54 | |
| Nucleotide | 104.45/274.02/156.85 | |
| Ligand | --- | |
| Water | --- | |
| **Data** | | |
| Resolution Estimates (Å) | Masked | Unmasked |
| d model | 3.8 | 3.7 |
| d FSC model (0/0.143/0.5) | 3.3/3.4/4.1 | 3.3/3.5/4.2 |
| Map min/max/mean | -45.1613 | --- |
| **Model vs. Data** | | |
| CC (mask) | 0.76 | |
| CC (box) | 0.85 | |
| CC (peaks) | 0.72 | |
| CC (volume) | 0.76 | |

**Supplementary Table 2. Data collection parameters and CryoEM map and model Validation and Refinement Statistics Using PHENIX**

| Protein | Amino acid sequence |
|---|---|
| **6xHis-MBP LbuCas13a** | MKSSHHHHHHGSSMKIEEGKLVIWINGDKGYNGLAEVGKKFEKDTGIKVTVEHPDKLEEKFPQVAATGD<br>GPDIIFWAHDRFGGYAQSGLLAEITPDKAFQDKLYPFTWDAVRYNGKLIAYPIAVEALSLIYNKDLLPNPPK<br>TWEEIPALDKELKAKGKSALMFNLQEPYFTWPLIAADGGYAFKYENGKYDIKDVGVDNAGAKAGLTFLVD<br>LIKNKHMNADTDYSIAEAAFNKGETAMTINGPWAWSNIDTSKVNYGVTVLPTFKGQPSKPFVGVLSAGIN<br>AASPNKELAKEFLENYLLTDEGLEAVNKDKPLGAVALKSYEEELAKDPRIAATMENAQKGEIMPNIPQMS<br>AFWYAVRTAVINAASGRQTVDEALKDAQTNSSSNNNNNNNNNNLGIEENLYFQSNAMKVTKVGGISHKK<br>YTSEGRLVKSESEENRTDERLSALLNMRLDMYIKNPSSTETKENQKRIGKLKKFFSNKMVYLKDNTLSLK<br>NGKKENIDREYSETDILESDVRDKKNFAVLKKIYLNENVNSEELEVFRNDIKKKLNKINSLKYSFEKNKANY<br>QKINENNIEKVEGKSKRNIIYDYYRESAKRDAYVSNVKEAFDKLYKEEDIAKLVLEIENLTKLEKYKIREFYH<br>EIIGRKNDKENFAKIIYEEIQNVNNMKELIEKVPDMSELKKSQVFYKYYLDKEELNDKNIKYAFCHFVEIEMS<br>QLLKNYVYKRLSNISNDKIKRIFEYQNLKKLIENKLLNKLDTYVRNCGKYNYYLQDGEIATSDFIARNRQNE<br>AFLRNIIGVSSVAYFSLRNILETENENDITGRMRGKTVKNNKGEEKYVSGEVDKIYNENKKNEVKENLKMF<br>YSYDFNMDNKNEIEDFFANIDEAISSIRHGIVHFNLELEGKDIFAFKNIAPSEISKKMFQNEINEKKLKLKIFR<br>QLNSANVFRYLEKYKILNYLKRTRFEFVNKNIPFVPSFTKLYSRIDDLKNSLGIYWKTPKTNDDNKTKEIIDA<br>QIYLLKNIYYGEFLNYFMSNNGNFFEISKEIIELNKNDKRNLKTGFYKLQKFEDIQEKIPKEYLANIQSLYMIN<br>AGNQDEEEKDTYIDFIQKIFLKGFMTYLANNGRLSLIYIGSDEETNTSLAEKKQEFDKFLKKYEQNNNIKIP<br>YEINEFLREIKLGNILKYTERLNMFYLILKLLNHKELTNLKGSLEKYQSANKEEAFSDQLELINLLNLDNNRV<br>TEDFELEADEIGKFLDFNGNKVKDNKELKKFDTNKIYFDGENIIKHRAFYNIKKYGMLNLLEKIADKAGYKIS<br>IEELKKYSNKKNEIEKNHKMQENLHRKYARPRKDEKFTDEDYESYKQAIENIEEYTHLKNKVEFNELNLLQ<br>GLLLRILHRLVGYTSIWERDLRFRLKGEFPENQYIEEIFNFENKKNVKYKGGQIVEKYIKFYKELHQNDEVK<br>INKYSSANIKVLKQEKKDLYIRNYIAHFNYIPHAEISLLEVLENLRKLLSYDRKLKNAVMKSVVDILKEYGFV<br>ATFKIGADKKIGIQTLESEKIVHLKNLKKKKLMTDRNSEELCKLVKIMFEYKMEEKKSEN |
| **6xHis-MBP LbuCas13a Δ409-421** | MKSSHHHHHHGSSMKIEEGKLVIWINGDKGYNGLAEVGKKFEKDTGIKVTVEHPDKLEEKFPQVAATG<br>DGPDIIFWAHDRFGGYAQSGLLAEITPDKAFQDKLYPFTWDAVRYNGKLIAYPIAVEALSLIYNKDLLPN<br>PPKTWEEIPALDKELKAKGKSALMFNLQEPYFTWPLIAADGGYAFKYENGKYDIKDVGVDNAGAKAGL<br>TFLVDLIKNKHMNADTDYSIAEAAFNKGETAMTINGPWAWSNIDTSKVNYGVTVLPTFKGQPSKPFVG<br>VLSAGINAASPNKELAKEFLENYLLTDEGLEAVNKDKPLGAVALKSYEEELAKDPRIAATMENAQKGEI<br>MPNIPQMSAFWYAVRTAVINAASGRQTVDEALKDAQTNSSSNNNNNNNNNNLGIEENLYFQSNAMKV<br>TKVGGISHKKYTSEGRLVKSESEENRTDERLSALLNMRLDMYIKNPSSTETKENQKRIGKLKKFFSNK<br>MVYLKDNTLSLKNGKKENIDREYSETDILESDVRDKKNFAVLKKIYLNENVNSEELEVFRNDIKKKLNKI<br>NSLKYSFEKNKANYQKINENNIEKVEGKSKRNIIYDYYRESAKRDAYVSNVKEAFDKLYKEEDIAKLVLEI<br>ENLTKLEKYKIREFYHEIIGRKNDKENFAKIIYEEIQNVNNMKELIEKVPDMSELKKSQVFYKYYLDKEEL<br>NDKNIKYAFCHFVEIEMSQLLKNYVYKRLSNISNDKIKRIFEYQNLKKLIENKLLNKLDTYVRNCGKYNYY<br>LQDGEIATSDFIARNRQNEAFLRNIIGVSSVAYFSLRNILETENENDITGRMRGSGEVDKIYNENKKNEV<br>KENLKMFYSYDFNMDNKNEIEDFFANIDEAISSIRHGIVHFNLELEGKDIFAFKNIAPSEISKKMFQNEINE<br>KKLKLKIFRQLNSANVFRYLEKYKILNYLKRTRFEFVNKNIPFVPSFTKLYSRIDDLKNSLGIYWKTPKTN<br>DDNKTKEIIDAQIYLLKNIYYGEFLNYFMSNNGNFFEISKEIIELNKNDKRNLKTGFYKLQKFEDIQEKIPK<br>EYLANIQSLYMINAGNQDEEEKDTYIDFIQKIFLKGFMTYLANNGRLSLIYIGSDEETNTSLAEKKQEFDK<br>FLKKYEQNNNIKIPYEINEFLREIKLGNILKYTERLNMFYLILKLLNHKELTNLKGSLEKYQSANKEEAFSD<br>QLELINLLNLDNNRVTEDFELEADEIGKFLDFNGNKVKDNKELKKFDTNKIYFDGENIIKHRAFYNIKKYG<br>MLNLLEKIADKAGYKISIEELKKYSNKKNEIEKNHKMQENLHRKYARPRKDEKFTDEDYESYKQAIENIE<br>EYTHLKNKVEFNELNLLQGLLLRILHRLVGYTSIWERDLRFRLKGEFPENQYIEEIFNFENKKNVKYKGG<br>QIVEKYIKFYKELHQNDEVKINKYSSANIKVLKQEKKDLYIRNYIAHFNYIPHAEISLLEVLENLRKLLSYD<br>RKLKNAVMKSVVDILKEYGFVATFKIGADKKIGIQTLESEKIVHLKNLKKKKLMTDRNSEELCKLVKIMFE<br>YKMEEKKSEN |
| **6xHis-MBP LbaCas13a** | MKSSHHHHHHGSSMKIEEGKLVIWINGDKGYNGLAEVGKKFEKDTGIKVTVEHPDKLEEKFPQVAATG<br>DGPDIIFWAHDRFGGYAQSGLLAEITPDKAFQDKLYPFTWDAVRYNGKLIAYPIAVEALSLIYNKDLLPN<br>PPKTWEEIPALDKELKAKGKSALMFNLQEPYFTWPLIAADGGYAFKYENGKYDIKDVGVDNAGAKAGL<br>TFLVDLIKNKHMNADTDYSIAEAAFNKGETAMTINGPWAWSNIDTSKVNYGVTVLPTFKGQPSKPFVG<br>VLSAGINAASPNKELAKEFLENYLLTDEGLEAVNKDKPLGAVALKSYEEELAKDPRIAATMENAQKGEI<br>MPNIPQMSAFWYAVRTAVINAASGRQTVDEALKDAQTNSSSNNNNNNNNNNLGIEENLYFQSNAMKI<br>SKVREENRGAKLTVNAKTAVVSENRSQEGILYNDPSRYGKSRKNDEDRDRYIESRLKSSGKLYRIFNE<br>DKNKRETDELQWFLSEIVKKINRRNGLVLSDMLSVDDRAFEKAFEKYAELSYTNRRNKVSGSPAFETC<br>GVDAATAERLKGIISETNFINRIKNNIDNKVSEDIIDRIIAKYLKKSLCRERVKRGLKKLLMNAFDLPYSDP<br>DIDVQRDFIDYVLEDFYHVRAKSQVSRSIKNMNMPVQPEGDGKFAITVSKGGTESGNKRSAEKEAFKK<br>FLSDYASLDERVRDDMLRRMRRLVVLYFYGSDDSKLSDVNEKFDVWEDHAARRVDNREFIKLPLENK<br>LANGKTDKDAERIRKNTVKELYRNQNIGCYRQAVKAVEEDNNGRYFDDKMLNMFFIHRIEYGVEKIYA<br>NLKQVTEFKARTGYLSEKIWKDLINYISIKYIAMGKAVYNYAMDELNASDKKEIELGKISEEYLSGISSFD<br>YELIKAEEMLQRETAVYVAFAARHLSSQTVELDSENSDFLLLKPKGTMDKNDKNKLASNNILNFLKDKE<br>TLRDTILQYFGGHSLWTDFPFDKYLAGGKDDVDFLTDLKDVIYSMRNDSFHYATENHNNGKWNKELIS<br>AMFEHETERMTVVMKDKFYSNNLPMFYKNDDLKKLLIDLYKDNVERASQVPSFNKVFVRKNFPALVR<br>DKDNLGIELDLKADADKGENELKFYNALYYMFKEIYYNAFLNDKNVRERFITKATKVADNYDRNKERNL<br>KDRIKSAGSDEKKKLREQLQNYIAENDFGQRIKNIVQVNPDYTLAQICQLIMTEYNQQNNGCMQKKSA<br>ARKDINKDSYQHYKMLLLVNLRKAFLEFIKENYAFVLKPYKHDLCDKADFVPDFAKYVKPYAGLISRVA<br>GSSELQKWYIVSRFLSPAQANHMLGFLHSYKQYVWDIYRRASETGTEINHSIAEDKIAGVDITDVDAVID<br>LSVKLCGTISSEISDYFKDDEVYAEYISSYLDFEYDGGNYKDSLNRFCNSDAVNDQKVALYYDGEHPKL<br>NRNIILSKLYGERRFLEKITDRVSRSDIVEYYKLKKETSQYQTKGIFDSEDEQKNIKKFQEMKNIVEFRDL |

| | |
|---|---|
| | MDYSEIADELQGQLINWIYLRERDLMNFQLGYHYACLNNDSNKQATYVTLDYQGKKNRKINGAILYQIC<br>AMYINGLPLYYVDKDSSEWTVSDGKESTGAKIGEFYRYAKSFENTSDCYASGLEIFENISEHDNITELR<br>NYIEHFRYYSSFDRSFLGIYSEVFDRFFTYDLKYRKNVPTILYNILLQHFVNVRFEFVSGKKMIGIDKKDR<br>KIAKEKECARITIREKNGVYSEQFTYKLKNGTVYVDARDKRYLQSIIRLLFYPEKVNMDEMIEVKEKKKP<br>SDNNTGKGYSKRDRQQDRKEYDKYKEKKKKEGNFLSGMGGNINWDEINAQLKN |
| **6xHis-SUMO TccCas13a** | MGSSHHHHHHSSGLVPRGSHMSGSAAGGEEDKKPAGGEGGGAHINLKVKGQDGNEVFFRIKRSTQ<br>LKKLMNAYCDRQSVDMTAIAFLFDGRRLRAEQTPDELEMEDGDEIDAMLHQTGGMKITKRKWGEHHP<br>PLYFYRDEDSGRLLAQNDRKQDYTDTLFNDIAQDTFERSLRNRLLKTPEKGDKRFYSNEIVKLVEKLC<br>QGADVAEIMKSMERNEKLRPKNEKEIKNLKKQLDGTLSEYGKRYTAPEGAMTLNDALFYLVEGNPLKQ<br>AMAKAELGKIREALIKEKENRINRVRYSIKNNKIPLRIQEDGGITPNNDRAAWLLGLMKPADPAKGITDC<br>YPLLGELEEVFDFDKLSKTLHEKISRCQGRPRSIAMAVDEALKQYLRELWEKSPSRQQDLKYYFQAVQ<br>EYFKDNFPIRTKRMGARLRQELLKDKTSLSRLLEPKHMANAVRRRLINQSTQMHILYGKLYAYCCGED<br>GRLLVNSETLQRIQVHEAVKKQAMTAVLWSISRLRYFYQFEDGDILSNKNPIKDFRDKFLRDTNKYTHE<br>DVEACKEKLQDFFPLKELQEKIKEDAKGLQETDNKQADTTDFKAIGHIVRDDRKLCNQLLAECVSCIGE<br>LRHHIFHYKNVTLIQALKRIADKVKPEDLSVLRAIYLLDRRNLKKAFAKRISSMNLPLYYREDLLSRIFKKE<br>GTAFFLYSAKIQMTPSFQRVYERGKNLRREFECERMKAEASNGQNGQDGDRLKWFRQLAAGDSADT<br>HFNWAVEAYAESAADVENNVEFDTDVDAQRALRNLLLLIYRHHFLPEVQKDETLVTGKIHKVLERNRQ<br>LSEGQGPNQGKAHGYSVIEELYHEGMPLSDLMKQLQRRISETERESRELAQEKTDYAQRFILDIFAEA<br>FNDFLEAHYGEEYLEIMSPRKDAEAAKKWVKESKTVDLKTSIDEKEPEGHLLVLYPVLRLLDERELGEL<br>QQQMIRYRTSLASWQGESNFSEEIRIAGQIEELTELVKLTEPEPQFAEEVWGKRAKEAFEDFIEGNMK<br>NYEAFYLQSDNNTPVYRRNMSRLLRSGLMGVYQKVLASHKQALKRDYLLWSEKHWNVKDENGADIS<br>SAEQAQCLLQRLHRKYAESPSRFTEEDCKLYEKVLRRLEDYNQAVKNLSFSSLYEICVLNLEILSRWVG<br>FVQDWERDMYFLLLAWVRQGKLDGIKEEDVRDIFSEGNIIRNLVDTLKGENMNAFESVYFPENKGSKY<br>LGVRNDVAHLDLMRKNGWRLEAGKTCSVMEDYINRLRFLLSYDQKRMNAVTKTLQQIFDRHKVKIRFT<br>VEKGGMLKIEDVTADKIVHLKGSRLSGIEIPSHGERFIDTLKALMVYPRG |
| **RfxCas13d-6xHis** | MIEKKKSFAKGMGVKSTLVSGSKVYMTTFAEGSDARLEKIVEGDSIRSVNEGEAFSAEMADKNAGYKI<br>GNAKFSHPKGYAVVANNPLYTGPVQQDMLGLKETLEKRYFGESADGNDNICIQVIHNILDIEKILAEYIT<br>NAAYAVNNISGLDKDIIGFGKFSTVYTYDEFKDPEHHRAAFNNNDKLINAIKAQYDEFDNFLDNPRLGYF<br>GQAFFSKEGRNYIINYGNECYDILALLSGLRHWVVHNNEEESRISRTWLYNLDKNLDNEYISTLNYLYD<br>RITNELTNSFSKNSAANVNYIAETLGINPAEFAEQYFRFSIMKEQKNLGFNITKLREVMLDRKDMSEIRK<br>NHKVFDSIRTKVYTMMDFVIYRYYIEEDAKVAAANKSLPDNEKSLSEKDIFVINLRGSFNDDQKDALYYD<br>EANRIWRKLENIMHNIKEFRGNKTREYKKKDAPRLPRILPAGRDVSAFSKLMYALTMFLDGKEINDLLTT<br>LINKFDNIQSFLKVMPLIGVNAKFVEEYAFFKDSAKIADELRLIKSFARMGEPIADARRAMYIDAIRILGTN<br>LSYDELKALADTFSLDENGNKLKKGKHGMRNFIINNVISNKRFHYLIRYGDPAHLHEIAKNEAVVKFVLG<br>RIADIQKKQGQNGKNQIDRYYETCIGKDKGKSVSEKVDALTKIITGMNYDQFDKKRSVIEDTGRENAER<br>EKFKKIISLYLTVIYHILKNIVNINARYVIGFHCVERDAQLYKEKGYDINLKKLEEKGFSSVTKLCAGIDETA<br>PDKRKDVEKEMAERAKESIDSLESANPKLYANYIKYSDEKKAEEFTRQINREKAKTALNAYLRNTKWN<br>VIIREDLLRIDNKTCTLFRNKAVHLEVARYVHAYINDIAEVNSYFQLYHYIMQRIIMNERYEKSSGKVSEY<br>FDAVNDEKKYNDRLLKLLCVPFGYCIPRFKNLSIEALFDRNEAAKFDKEKKKVSGNSSRGSHHHHHH |
| **AlcrVIA1-6xHis** | MKTIKVDVIVVGDDEELVEEYKKEAELIGKEYGVKIEVEPYFLEEGKFPWLDVDFAYNTTQEELDKAEK<br>EAKKIAGSHHHHHH |
| **AlcrVIA2-6xHis** | MNEKKEKARKEMKEIAEKAIKEIKKDPEKALEIAVKAVEEIGEIANESGDYKSGEENAWKVAEEYAKVT<br>GDDDTAYDLMLAGDMIIDGKDAEEGYEILKELGSHHHHHH |
| **AlcrVIA3-6xHIs** | MAPKKYVVTVTIPVTDADLTVFLVVDIIVYAEKLGGTVTITAVKSENDSYSVTLEDLDKAAEELEKVGGSI<br>VLTVTFDNKEKAEKVAEFAYLKAEEYNLKVDVEVKEELGSHHHHHH |

**Supplementary Table 3. Cas13 and AlcrVIA Sequences**

| **Name** | **Sequence (5′- 3′)** | **Notes** |
|---|---|---|
| **T7 FOR primer** | GCGCGCATTAATACGACTCACTATAGGG | |
| **T7 REV primer** | CAAAAAACCCCTCAAGAC | |
| **LbuCas13a crRNA** | UAGACCAGCCCAAAAAUGAAGGGCACUAAAACGCAGCGCCUCUUGCAACGAUUAAA | |
| **LbaCas13a crRNA** | AGAAGAUAGCCCAAGAAAGAGGGCAAUAACGCAGCGCCUCUUGCAACGAUUAAA | |
| **TcaCas13a crRNA** | CACAACUCCCAUGUAGGCGGAGACUGCAACGCAGCGCCUCUUGCAACGAUUAAG | |
| **RfxCas13a crRNA** | AACCCCUACCAACUGGUCGGGGUUUGAAACGCAGCGCCUCUUGCAACGAUUAAA | |
| **Cas13 aRNA** | UUUAAUCGUUGCAAGAGGCGCUGCUC | |
| **PolyU reporter** | 56-FAM UUUUU IowaBlck | |
| **PolyA reporter** | 56-FAM AAAAA IowaBlck | |
| **PolyCG reporter** | 56-FAM CGCGC IowaBlck | |
| **Fluorescence polarisation aRNA** | 56-FAM C*CGGCAGAAAAAGGAAAGAAAGAAACC | **C*= deoxyribose** |
| **cRNA cryoEM and BLI experiment** | UAGACCACCCCAAAAAUGAAGGGGACUAAAACUUUCUUUCUUUCCUUUUUCUGCCG | |
| **aRNA cryoEM and BLI experiment** | CGGCAGAAAAAGGAAAGAAAGAAACC | |

**Supplementary Table 4. List of DNA and RNA oligonucleotides**
