## Supplementary figures and images for "*De novo* design of potent CRISPR-Cas13 inhibitors"

### Supplementary Video 1

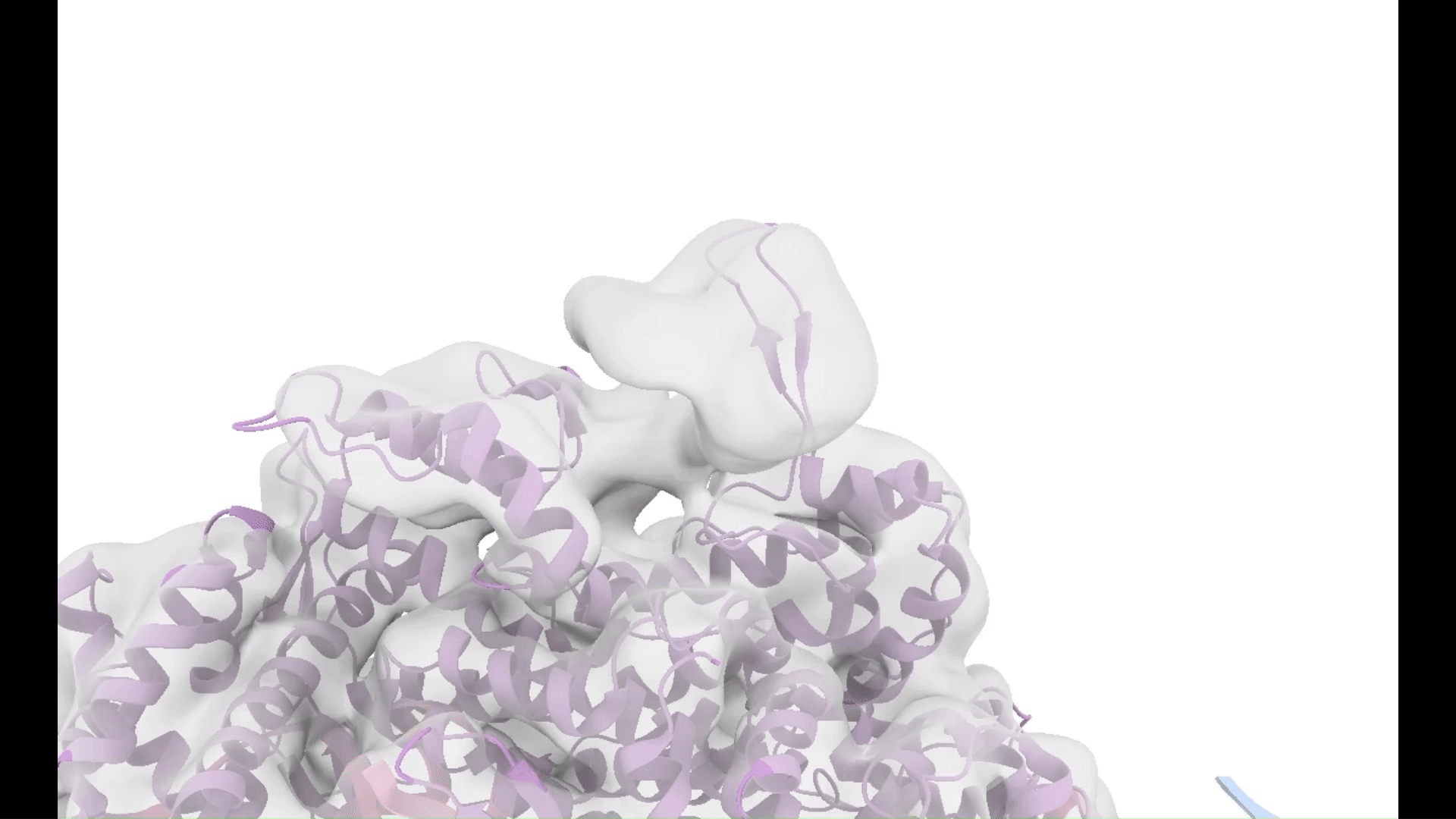
